# Disease-associated gut microbiome and metabolome changes in chronic low back pain patients with bone marrow lesions

**DOI:** 10.1101/2023.07.26.550629

**Authors:** Wentian Li, Ji Tu, Jinjian Zheng, Abhirup Das, Qi Yan, Xiaotao Jiang, Wenyuan Ding, Xupeng Bai, Kaitao Lai, Sidong Yang, Cao Yang, Jun Zou, Ashish D Diwan, Zhaomin Zheng

**Affiliations:** Spine Labs, St. George and Sutherland Clinical School, University of New South Wales, Kogarah, NSW 2217, Australia; Department of Spine Surgery, The First Affiliated Hospital, Sun Yat-Sen University, Guangzhou, China; Department of Orthopedic Surgery, The First Affiliated Hospital of Soochow University, Suzhou, Jiangsu 215006, China; UNSW Microbiome Research Centre, St George and Sutherland Clinical Campuses, School of Clinical Medicine, UNSW Medicine and Health, The University of New South Wales, Sydney, NSW 2052, Australia; Department of Spinal Surgery, The Third Hospital of Hebei Medical University, 139 Ziqiang Road, Shijiazhuang 050051, China; Hebei Joint International Research Centre for Spinal Diseases; Center for Innovation & Translational Medicine, the First Affiliated Hospital, Zhejiang University School of Medicine, Hangzhou, China; Zhejiang Provincial Key Laboratory of Pancreatic Disease, the First Affiliated Hospital, Zhejiang University School of Medicine, Hangzhou, China; Charles Perkins Centre and School of Medical Sciences, University of Sydney, Sydney, Australia; ANZAC Research Institute, Concord Hospital, Sydney, Australia; Tissue Engineering and Microfluidics Laboratory (TE&M), Australian Institute for Bioengineering and Nanotechnology (AIBN), The University of Queensland, St Lucia, 4072, Queensland, Australia; Department of Orthopedics, Union Hospital, Tongji Medical College, Huazhong University of Science and Technology, Wuhan 430022, China; Spine Service, Department of Orthopedic Surgery, St. George Hospital, Kogarah, NSW 2217, Australia

**Keywords:** Chronic low back pain, Bone marrow lesions, Fatty replacement, Gut microbiome, Serum metabolomics, BM-MSCs

## Abstract

Chronic low back pain (LBP) is the leading cause of global disability. Vertebral bone marrow lesions (BMLs), one etiological factor for chronic LBP, are MRI signal changes in the vertebral bone marrow that extend from the disc endplate. The adipogenesis of bone marrow mesenchymal stem cells (BM-MSCs) could explain fatty replacement (FR) in normal bone marrow. FR is the most common type of BMLs. Here we show how the gut microbiome and serum metabolome change and how they interact in LBP patients with or without FR. The serum metabolome of chronic LBP patients with FR is characterized by decreased levels of branched-chain amino acids (BCAAs), which correlate with a gut microbiome that has important capability to regulate BCAA degradation pathway. *Ruminococcus gnavus, Roseburia hominis and Lachnospiraceae bacterium 8 1 57FAA* are identified as the main species driving the association between biosynthesis of BCAAs and BM-MSCs metabolism in LBP with FR individuals. *In vitro* work demonstrates that BCAAs can induce the adipogenesis of BM-MSCs by activating the SIRT4 pathway. Our findings provide a deep insight into understanding the role of the disturbed gut ecosystem in FR and LBP.

## Introduction

Chronic low back pain (LBP) is a common and debilitating condition worldwide, with the Lancet group calling for action^1^. Although chronic LBP relates to different spinal pathologies, vertebral bone marrow lesions (BMLs) on magnetic resonance imaging (MRI) have a high specificity for discogenic LBP^2^. Vertebral BMLs are pathological changes in the bone marrow composition in vertebral bodies. A meta-analysis by Jensen et al. established that the prevalence of BMLs and the association with chronic LBP estimated the median prevalence of LBP types in symptomatic populations at 43% compared to only 6% in non-symptomatic populations^3^. Fatty replacement (FR) of normal bone marrow is the most frequent in vertebral BML appearances accounting for up to 90% of BMLs observed^4, 5^. Currently, the pathogenesis of vertebral BMLs remains unclear. FR happens in the vertebral bodies, specifically, in the bone marrow adjacent to the endplate. (Supplementary figure 1)

**Supplementary figure 1:**
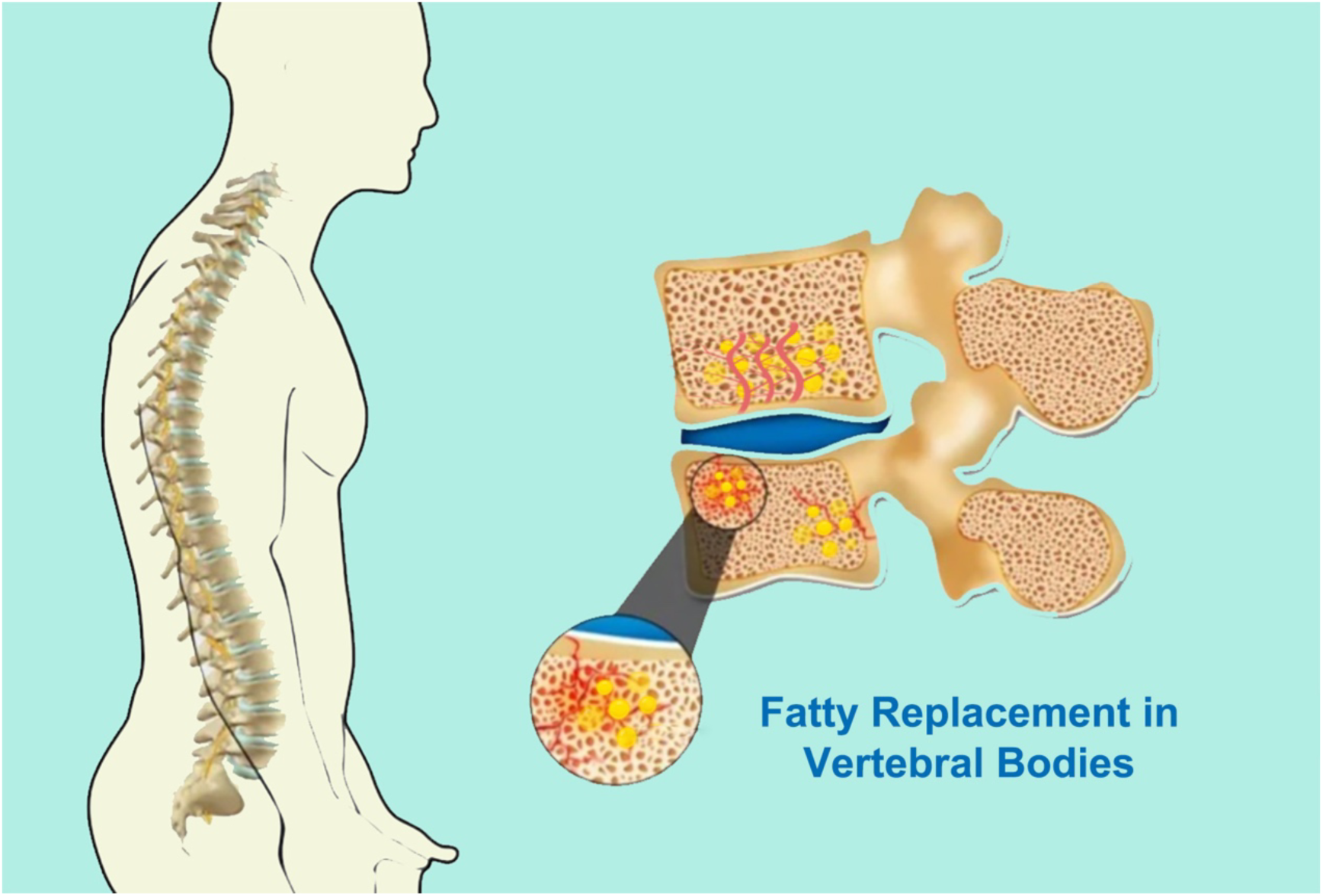
Illustration of chronic low back pain with fatty replacement.

The bone marrow microenvironment is filled with mesenchymal stem cells (BM- MSCs). BM-MSCs are multipotent stem cells that can differentiate into several mature cells, such as adipocytes and osteoblasts. As common progenitor cells of adipocytes and osteoblasts, BM-MSCs have delicately balanced the differentiation of osteogenesis and adipogenesis in the bone marrow. ^6^ Chronic LBP with FR was visualized as a fat replacement in vertebral bone marrow on MRI.^7^ Metabolism is vitally significant in BM-MSCs’ fate determination and differentiation.^8^ Thus, we hypothesize that chronic LBP with FR result from dysfunction of BM-MSC differentiation regulated by cellular metabolism.

Our understanding of the microbiome that lives in our gut, its functions, and its roles in human health and disease has advanced significantly over the last decade, aided by rapid technological advancement. Gut-bone axis was proposed to play a significant role in the onset of several bone-joint diseases such as osteoporosis, rheumatoid arthritis and spinal cord multiple sclerosis^9^. Previous studies indicated that gut microbiome communicated with bone marrow, regulating disease pathogenesis^10^. As the FR are alterations in the bone marrow milieu in patients with chronic LBP, such as increased bone marrow adiposity, whether the gut microbiome impacts these alterations is still unknown.

Longstreth and Yao drew attention to an excess of back surgery in patients with irritable bowel syndrome (IBS).^11, 12^ It has been shown that back pain is associated with altered gut microbiota composition. ^13^Recent research showed that LBP with BMLs could be caused by one specific microbiota (formerly known as *Propionibacterium acnes*, *P. acnes*). ^14^ This type of bacteria is an aerotolerant Gram-positive and a common skin commensal. Several animal models have found that *P. acnes* could exist in the degenerated discs and cause BMLs^15^. Considering the close relationship between LBP and BMLs, it is rational to hypothesize the potential roles of human microbiome in BMLs. However, thus far, no research has been done to detect whether gut microbiome disturbance accompanies BMLs.

There might be three potential mechanisms by which the gut microbiota could induce intervertebral disc (IVD) degeneration and cause chronic LBP with BMLs:(a) bacteria translocate across the gut epithelial barrier and arrive at IVDs or bone marrow. IVDs provide an excellent environment for anaerobic bacteria growth because of the low oxygen tension and the absence of immune surveillance; (b) translocation of the bacteria could regulate the mucosal and systemic immune systems; (c) the nutrients and metabolites formatted in the gut epithelium diffuse into the IVDs or bone marrow, and then cause FR.^16^

The human gut microbiota produces numerous metabolites. These metabolites could be accumulated in the bloodstream, which can have systemic effects on the host^17^. Serum levels of amino acids, most consistently the BCAAs^18^, were the most consumed metabolites in BM-MSCs differentiated into adipocytes. Interestingly, mature adipocytes have been shown to utilize BCAAs for acetyl-coenzyme A (CoA) production for lipogenesis^19^. However, the correlation between the gut microbiome and serum metabolome in LBP patients is unknown.

To bridge the abovementioned gaps, we conducted 16S and shotgun metagenomics analysis of fecal samples from 107 chronic LBP patients with or without FR and 31 healthy volunteers to define their composition of gut microbiota. We first combined shotgun metagenomics and metabolic analysis to uncover the gut microbiome’s functional and taxonomic characteristics in chronic LBP patients with or without FR and healthy volunteers. After that, we evaluated microbiomes’ influence on the capabilities of BM-MSC differentiation. By integrating these multilevel omics results, we defined and characterized the different gut microbiome and metabolites and their combination and interaction in the gut-bone marrow ecosystem of chronic LBP with FR.

## Results

### 1. Clinical characteristics of the participants

We recruited 107 chronic LBP patients and 31 healthy controls (HC). In 107 LBP patients, 54 participants had LBP with FA, and 53 patients were LBP (LBP without FR). Hence, we focused on the microbiome analysis of LBP+FR, LBP (LBP without FR), and HC. There was no significant difference in gender, age, body mass index (BMI), total Cholesterol Levels, and Triglycerides Levels between the three groups. We found that the LBP+FR cohort showed significantly higher Visual Analog Scale (VAS) and Oswestry Disability Index (ODI) scores than the LBP and HC cohorts. (Table 1) These well-matched samples were used to identify molecular signatures in the gut microbiome that modulate host metabolism.

**Table 1.** Characteristics of the study population.

|  | **LBP+FR** | **LBP** | **HC** | **P-value** |
| --- | --- | --- | --- | --- |
|  | **(n= 54)** | **(n= 53)** | **(n= 31)** |  |
| **Age (years)** | 45.67 ± 6.27 | 45.91 ± 8.72 | 42.10 ± 9.50 | Ns |
| **Gender (F/M)** | 26 / 28 | 29 / 24 | 13 / 18 | Ns |
| **BMI, mean (Kg/m<sup>2</sup>)** | 22.20 ± 5.01 | 23.83 ± 2.44 | 23.10 ± 2.10 | Ns |
| **VAS (1-10)** | 5.31 ± 1.84 | 4.43 ± 2.03 | ~ | 0.02 |
| **ODI (%)** | 46.2 ± 19.1% | 38.5 ± 17.7% | ~ | 0.03 |
| **Total cholesterol level (mmol/L)** | 4.55 ± 1.08 | 4.42 ± 1.04 | 4.47 ± 0.71 | Ns |
| **Total triglycerides level (mmol/L)** | 1.40 ± 0.72 | 1.43 ± 0.89 | 1.45 ± 0.72 | Ns |
| **Diet** | Ad libitum diet | Ad libitum diet | Ad libitum diet |  |

Values are presented as mean ± standard deviation. HC = Healthy Controls; LBP+FR
= Low back Pain with Fatty Replacement; LBP = Low Back Pain without Fatty
Replacement; BMI = Body Mass Index; VAS = Visual Analog Scale for pain; ODI =
Oswestry Disability Index.

### 2. Fecal microbiome taxonomic indicators of LBP with FR using 16S rRNA gene sequencing

To compare the gut bacterial community composition between LBP with FA, LBP patients and healthy controls, we first performed 16S rRNA sequencing for all 138 fecal samples. 16S rRNA sequencing confirmed a dysbiosis in LBP+FR and LBP compared to HC (Fig. 1A, B). Initially, alpha-diversity analysis of gut microbiome indicated samples from LBP+FR and LBP had decreased alpha-diversity index: Chao-1-richness index (Fig. 1A), Observed-otus, Shannon’s diversity index and PD-whole-tree index (Supplementary Fig. 2A-C), compared with samples from HC group. Besides, bacterial alpha-diversity analysis showed no significant difference among these indexes between LBP+FR and LBP groups.

**Figure 1.**
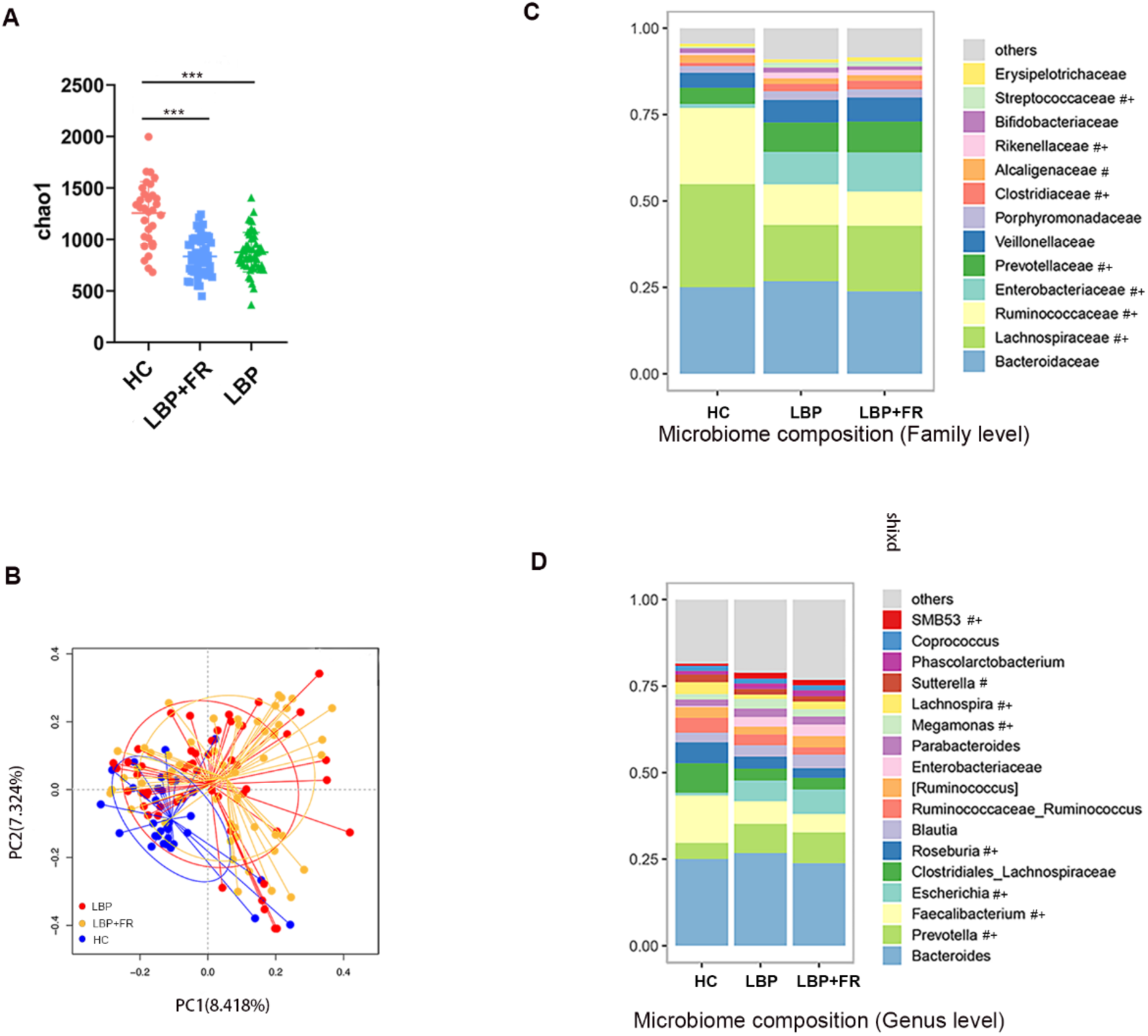
Distinct fecal microbiota profiles of subjects with LBP+FR, LBP and HC by using 16S rRNA gene amplicon sequencing. (A) Comparison of alpha-diversity indices (Chao-1-richness index) between HC, LBP+FR and LBP groups. (B) Principal coordinate analysis (PCoA) based on Bray Curtis distances demonstrating the separation of HC, LBP+FR and LBP groups. (C) Microbiome composition at family level in HC, LBP+FR and LBP fecal samples. (D) Microbiome composition at genus level showing the top 17 most abundant genus HC, LBP+FR and LBP fecal samples. Data in C-D are presented as mean relative abundance, with differences between groups shown as #*P* < 0.05 for LBP+FR compared to HC control, and +*P* < 0.05 for LBP compared to HC control, with exact *P* values shown in Supplementary Table 2.

Next, β-diversity analysis was used to explore patients’ overall gut microbiome phenotypes between three patient groups. Principal coordinate analysis (PCoA) showed that bacterial signatures between LBP+FR and LBP when compared to HC group were significantly distinct when using the Bray_Curtis distance (*P* = 0.01) (Fig. 2B), Unweighted_UniFrac and Weighted_UniFrac distances (Supplementary Fig. 2D-E).

**Figure 2.**
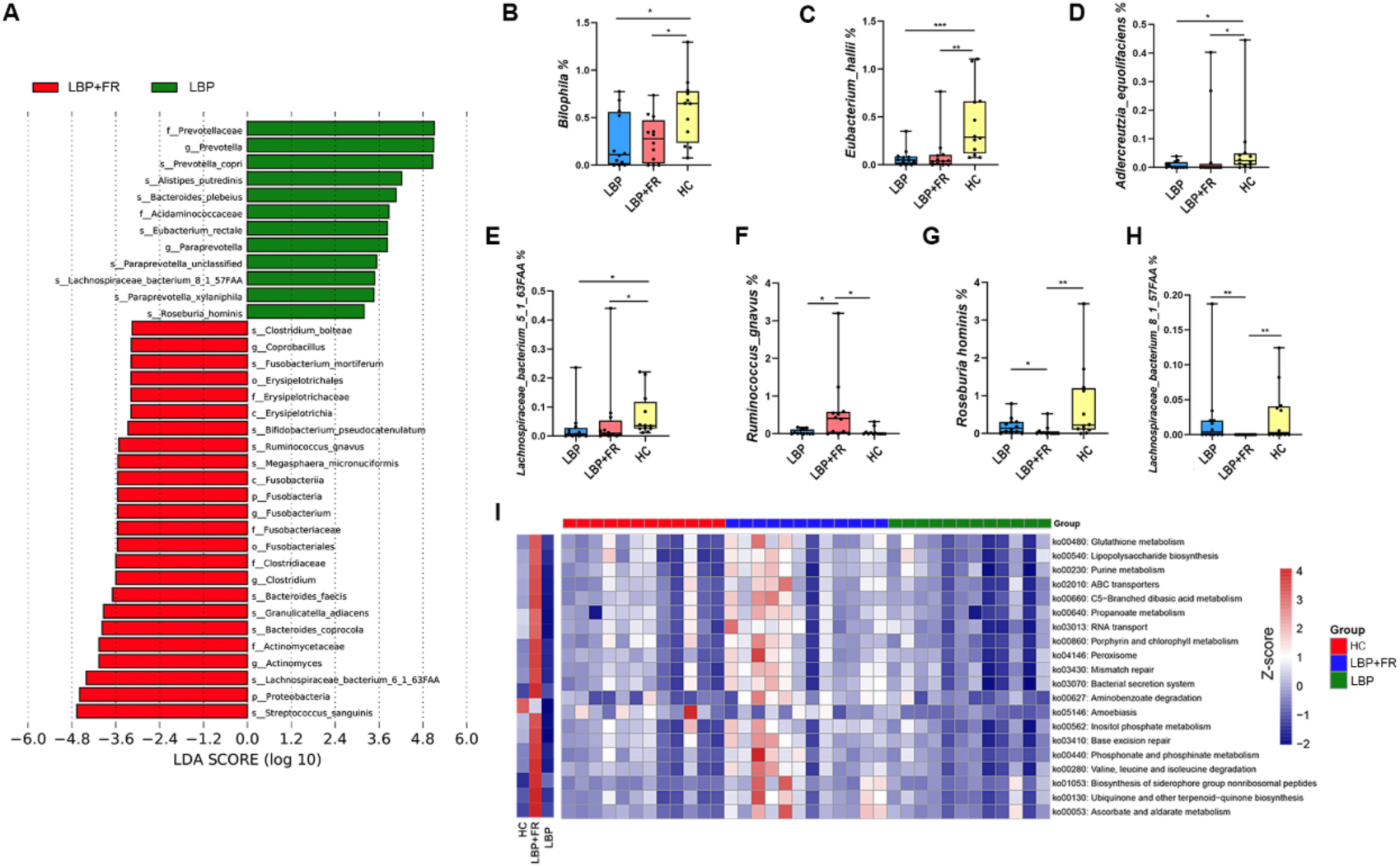
The fecal microbiota of LBP+FR patients can be distinguished from that of LBP and HC individuals in metagenomic analysis. (A) Linear discriminant analysis effect size identified the most differentially abundant taxa between the LBP+FR and LBP groups. LBP-enriched taxa are indicated with a positive LDA score, and taxa enriched in LBP+FR groups have a negative score. Only taxa meeting an LDA significant threshold of >2 are shown. (B-H) The relative abundances at species level of *Bilophila, Eubacterium hallii, Adlercreutzia equolifaciens, Lachnospiraceae bacterium 5 1 63FAA, Ruminococcus gnavus, Roseburia hominis* and *Lachnospiraceae bacterium 8 1 57FAA* between LBP+FR, LBP and HC groups. * indicates *p* <0.05; ** indicates *p* <0.01; and *** indicates *p* <0.001 by Wilcoxon rank-sum test. (I) Heatmap of functional capacity profiles showed the top 20 enrichments in the LBP+FR group by metagenomic sequencing analysis.

Here, we identified 343 discriminative bacterial species between three groups (Supplementary Table 1). Compared with HC cohort, LBP+FR and LBP cohorts were characterized by presence of five enriched families (Streptococcaceae, Prevotellaceae, Rikenellaceae, Clostridiaceae and Enterobacteriaceae) and by three depleted families (Ruminococcaceae, Alcaligenaceae and Lachnospiraceae) (Fig. 1C). No other differences were observed between the LBP with FR and LBP groups at the family level. Further, at the genus level, compared with HC, LBP+FR and LBP were confirmed by four enriched genuses (Prevotella, Escherichia, Megamonas and SMB53) and four depleted genuses (Fecalibacterium, Roseburia, Lachnospira and Sutterella) (Fig. 1D).

### 3. Fecal microbiome taxonomic indicators of LBP with FR using metagenomics

Having identified distinct FR-associated fecal taxa using 16S rRNA gene sequencing, we sought to increase the resolution of these findings via metagenomic sequencing. Only a portion of the gut microbiota population identified by shotgun sequencing could be detected by 16S rRNA gene sequencing. Shotgun sequencing out-performs 16S sequencing in identifying fewer common species when a significant number of reads are available. We chose a two-stage design (16S, first-stage; metagenomics, second- stage) in this study which is recommended by previous researchers.^20, 21^ 36 samples from 3 groups (LBP+FR, LBP and HC) were chosen for metagenomic sequencing. Alpha diversity analysis showed that there was no significant difference among these indexes (Observed-otus, Chao-1-richness index and Shannon’s diversity index) between any two groups (Supplementary Fig. 3A). PCoA based on Jaccard dissimilarity distances showed a significant difference between LBP+FR and LBP compared to HC controls (*P* = 0.002) (Supplementary Fig. 3B).

To further identify which bacterial taxa were distinct between the LBP+FR and LBP, we performed the linear discriminant analysis of effect size (LEfSe analysis). We identified 11 genera showing significant differences at the species level (Fig. 2A and Supplementary Fig. 3C-M). The abundance comparisons of predominant genera showed that *Fusobacterium mortiferum, Ruminococcus gnavus, Granulicatella adiacens* and *Streptococcus sanguinis* were significantly enriched, whereas *Bacteroides coprocola, Prevotella copri, Alistipes putredinis, Bacteroides plebeius, Paraprevotella, Lachnospiraceae bacterium 8 1 57FAA* and *Roseburia hominis* were depleted in patients with LBP+FR compared to LBP group (Supplementary Fig. 3C- M).

Moreover, examination of the microbiome at the species level identified a consortium of common bacterial species significantly depleted in both LBP+FR and LBP compared to HC groups (Fig. 2B-H). These included *Bilophila unclassified, Eubacterium hallii, Adlercreutzia equolifaciens* and *Lachnospiraceae bacterium 5 1 63FAA* (Fig. 2B-E). Furthermore, three species were found to discriminate LBP+FR from LBP and HC: enriched *Ruminococcus gnavus*, depleted *Roseburia hominis* and reduced *Lachnospiraceae bacterium 8 1 57FAA* (Fig. 2F-H). Enriched *Ruminococcus gnavus* was also seen at LEfSe analysis between three groups (Supplementary Fig. 3N), thereby confirming that *Ruminococcus gnavus* is specifically enriched in LBP+FR compared to LBP and HC controls (LDA log10 3.5).

We also compared the functionality of the fecal Microbiome in LBP+FR, LBP and HC groups based on metagenomics sequencing. We confirmed the twenty most abundant KO (KEGG ORTHOLOGY) pathways in LBP+FR group (Fig. 2I). The top five pathways were associated with ascorbate and aldarate metabolism (ko00053), ubiquinone and other terpenoid-quinone biosynthesis (ko00130), biosynthesis of siderophore group nonribosomal peptides (ko01053), valine, leucine and isoleucine degradation (ko00280) and phosphonate and phosphinate metabolism (ko00440) (Fig. 2I and Supplementary Table 3).

### 4. Functional indicators of the LBP+FR fecal metabolome

To find the microbe-host interaction in chronic LBP with FR, we collected 120 serum samples from 138 participants. We tried to discriminate the metabolic profiles between LBP+FR (n = 40), LBP (n = 40) and HC (n = 30). The overwhelming mass of research supports that gut microbiome could produce some end products of fermentation. These products may enter our circulation system by blood and influence our physiology. The serum metabolome can provide the functional readout of the gut microbiome.^22^

In this study, principal component analysis (PCA) revealed a significant but incomplete separation of LBP+FR and LBP (PERMANOVA, *P* = 0.001; Fig. 3A). Moreover, PCA also revealed incomplete separation between LBP+FR, LBP and HC (Supplementary Fig. 4). In LBP+FR and LBP groups, we detected a total of 745 biochemicals that differed significantly with importance in project (VIP) >1. The LBP+FR group showed depletion in 185 metabolites and enrichment in 560 metabolites compared to the LBP group. Of the top fifty indicator metabolites with the highest variable VIP score separating LBP+FR from LBP samples, these altered metabolites were mainly involved in Amino acid metabolism, Carbohydrate metabolism, Fatty Acyls metabolism, Glycerophospholipids metabolism, and Sphingolipids metabolism. (Fig. 3B) Furthermore, based on the KEGG database, we conducted a metabolic pathway analysis to help us understand the major biochemical metabolic pathways and signal transduction pathways involved in metabolites. The result showed that patients with LBP+FR were mainly characterized by disturbances of Valine, leucine and isoleucine degradation (BCAAs degradation), Sphingolipid metabolism, Porphyrin and chlorophyll metabolism, Nicotinate and nicotinamide metabolism, Neomycin, kanamycin and gentamicin biosynthesis, Drug metabolism - cytochrome P450, Amino sugar and nucleotide sugar metabolism. (Fig. 3C and Supplementary Table 4) Among these metabolisms, BCAAs degradation was the most significant (*P* = 0.0156). Then, we measured the absolute qualification of BCAAs in serum samples. The level of BCAA in LBP+FR decreased significantly compared with LBP and HC groups. (Fig. 3D) Integration of these findings showed that disturbance of BCAAs metabolism was particularly relevant to LBP+FR.

**Figure 3.**
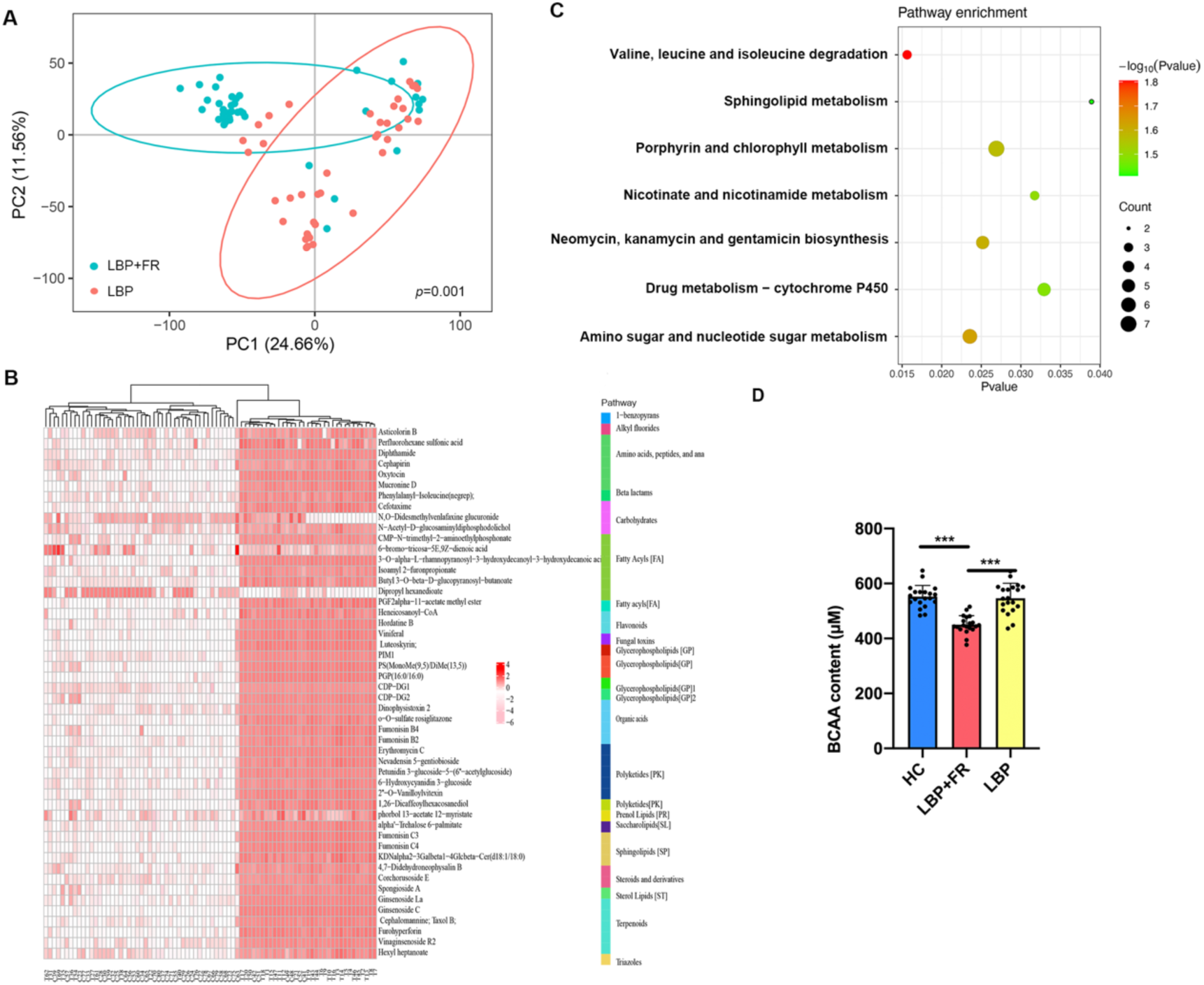
Blood metabolites that discriminate HC, LBP+FR and LBP. (A) Metabolic signatures of LBP+FR subjects were significantly distinguished from LBP (PERMANOVA, *P* = 0.001). Discovery set: LBP+FR, n = 40; LBP, n = 40. (B) Relative abundances of 50 blood metabolites differentiating between the two groups. Compared with LBP, the LBP+FR group was characterized by 16 up-regulated and 34 down-regulated metabolites. These metabolites were mainly involved in Amino acid metabolism, Carbohydrate metabolism, Fatty Acyls metabolism, Glycerophospholipids metabolism, and Sphingolipids metabolism. (C) Metabolomics pathway enrichment analysis identified seven significantly over- represented sub-pathways among the differential metabolites. (D) The absolute qualification of BCAAs in serum samples between LBP+FR (n = 20), LBP (n = 20) and HC groups (n = 20). * indicates *p* <0.0 5; ** indicates *p* <0.01; and *** indicates *p* <0.001 by Wilcoxon rank-sum test.

### 5. Co-occurrence analysis among the gut microbiome and metabolites

Co-occurrence analysis was used to explore the potential relationships between different microbiome species and serum metabolites. The analysis showed that the bacterial species formed strong and broad co-occurring relationships with serum metabolites. (Fig. 4)

**Figure 4.**
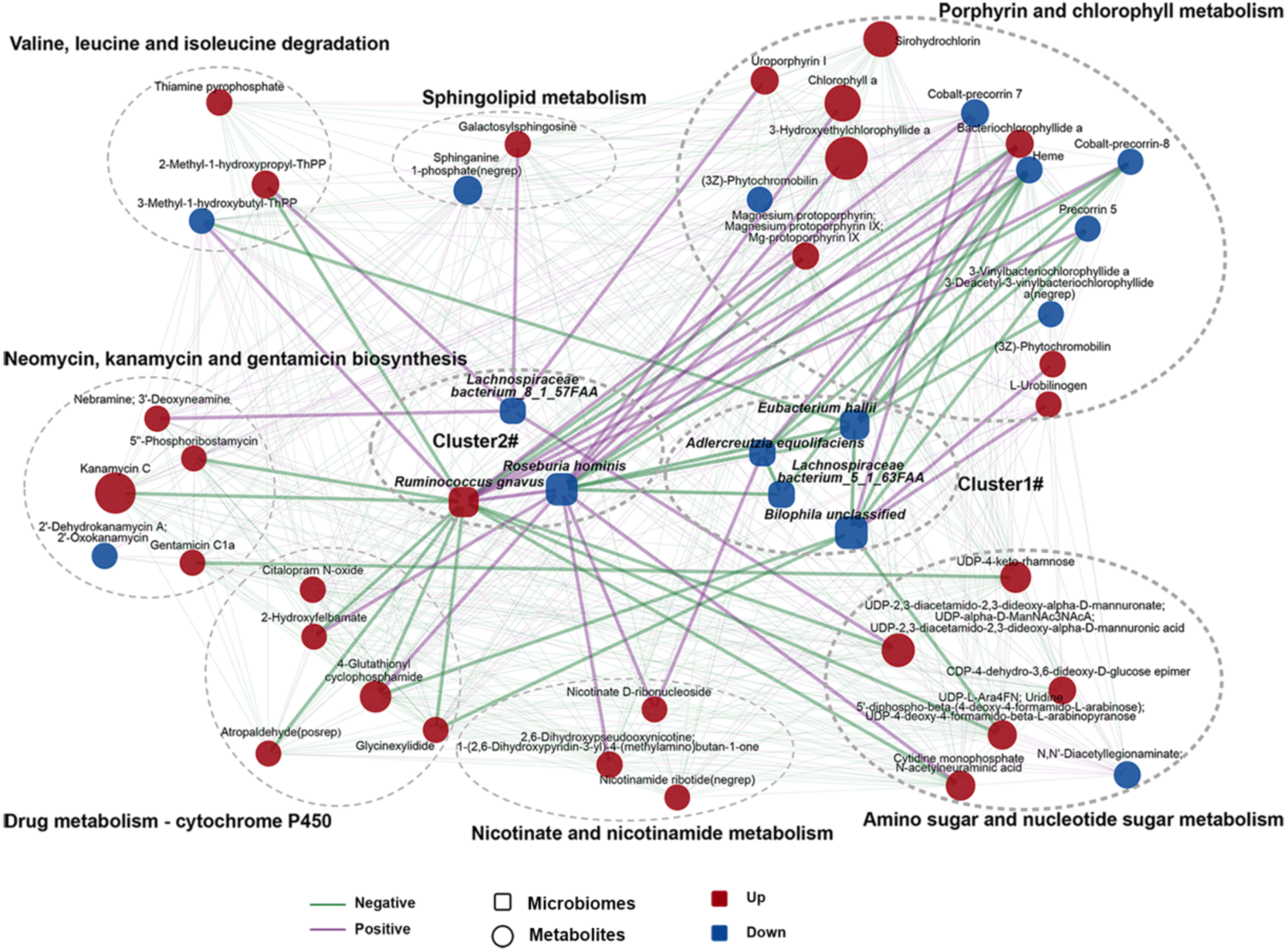
A co-occurrence network constructed from the relative abundances of differential bacteria, and serum metabolites in LBP+FR/LBP subjects versus HCs.

Within this co-expression network, these differential bacterial species were separated into two clusters (clusters 1 and 2). In the LBP with or without FR group relative to the HC group, cluster 1 was composed of four depleted bacterial species; (*Bilophila unclassified, Eubacterium hallii, Adlercreutzia equolifaciens* and *Lachnospiraceae bacterium 5 1 63FAA*). Cluster 2 mainly included three species (*Ruminococcus gnavus*, *Roseburia hominis* and *Lachnospiraceae bacterium 8 1 57FAA*) to distinguish LBP+FR group from LBP and HC groups. Within cluster 1, *Bilophila unclassified, Eubacterium hallii, Adlercreutzia equolifaciens* and *Lachnospiraceae bacterium 5 1 63FAA* showed negative correlation with each other, except for the connection between *Bilophila unclassified* and *Lachnospiraceae bacterium 5 1 63FAA*. Clusters 1 and 2 were linked by a common node (*Roseburia hominis*). In cluster 2, *Roseburia hominis* positively correlated with *Ruminococcus gnavus*. Besides, some members from cluster 1 (*Eubacterium hallii, Adlercreutzia equolifaciens* and *Lachnospiraceae bacterium 5 1 63FAA*) showed negative correlations with the members from cluster 2 (*Roseburia hominis*). These findings suggest that those important differential bacterial species may form a synergistic relationship in patients with LBP, LBP+FR and HC.

Regarding the correlation between microbiome and metabolites, the first network indicated associations between cluster 1 and a group of 14 metabolites in Porphyrin and chlorophyll metabolism. In our multivariate analysis, each of these 14 metabolites was identified as different between LBP (with/without FR) and HC group. The second network showed the connection between cluster 2 and numerous metabolites from 7 enriched pathways (Valine, leucine and isoleucine degradation, Sphingolipid metabolism, Porphyrin and chlorophyll metabolism, Nicotinate and nicotinamide metabolism, Neomycin, kanamycin and gentamicin biosynthesis, Drug metabolism - cytochrome P450 and Amino sugar and nucleotide sugar metabolism) (Fig. 3C). Significantly, Porphyrin and chlorophyll metabolism showed a strong correlation with both the cluster 1 species and cluster 2 species, which helps explain the close relationship of this pathway between LBP and related species. Moreover, the metabolites from the most significantly enriched pathway: Valine, leucine and isoleucine degradation (Fig. 3C), showed a positive correlation with *Ruminococcus gnavus* and *Lachnospiraceae bacterium 8 1 57FAA*, and a negative connection with *Ruminococcus gnavus*. Meanwhile, in cluster 2, *Ruminococcus gnavus* showed a more negative relationship with other metabolites, but *Roseburia hominis* and *Lachnospiraceae bacterium 8 1 57FAA* present a more positive connection with other metabolites. It suggested the different functions of these three species in the development of FR. Although our study found a potential interaction of gut microbiome with metabolites in LBP+FR, it remains to be further determined whether these metabolic products can influence the LBP+FR pathogenesis directly.

The discriminating bacterial species are mainly separated into two clusters. Cluster 1 was composed of 4 depleted species (*Bilophila unclassified, Eubacterium hallii,*

*Adlercreutzia equolifaciens* and *Lachnospiraceae bacterium 5 1 63FAA*) in LBP (with/without FR) subjects compared to HC group. Cluster 2 comprised three species (increased *Ruminococcus gnavus*, reduced *Roseburia hominis* and decreased *Lachnospiraceae bacterium 8 1 57FAA*). Within cluster 1, *Bilophila unclassified*, *Eubacterium hallii, Adlercreutzia equolifaciens* and *Lachnospiraceae bacterium 5 1 63FAA* showed negative correlation with each other, except the connection between *Bilophila unclassified* and *Lachnospiraceae bacterium 5 1 63FAA*. In cluster 2, *Ruminococcus gnavus* positively correlated with *Roseburia hominis*. Besides, some members from cluster 1 (*Eubacterium hallii, Adlercreutzia equolifaciens* and *Lachnospiraceae bacterium 5 1 63FAA)* showed negative correlations with the members from cluster 2 (*Roseburia hominis*).

In this network, altered metabolites in clusters 1 and 2 were mainly involved in Porphyrin and chlorophyll metabolism. Valine, leucine and isoleucine degradation positively correlated with *Ruminococcus gnavus* and *Lachnospiraceae bacterium 8 1 57FAA*, and a negative connection with *Ruminococcus gnavus*. The size of the nodes represents the abundance of these variables. Red and blue dots indicate the increased and decreased relative abundances of variables in LBP+FR/LBP subject relative to HC. Edges between nodes indicate Spearman’s negative (light green) or positive (light purse) correlation.

### 6. BM-MSCs from patients with LBP+FR exhibit a strong adipogenesis capability

After characterizing the gut microbiome composition and related metabolomics in LBP+FR and LBP, we tried to detect the effects of the gut microbiota on the BM-MSCs’ function in an ex vivo cell culture. We isolated in vitro BM-MSCs from the control, LBP+FR patients and LBP patients. Firstly, BM-MSCs were immune-phenotypically characterized by flow cytometry for the expression of mesenchymal, hematopoietic and neuronal markers. BM-MSCs were positive for CD105 and CD29, negative for CD34 and CD45 (Supplementary Fig. 5).

We stimulated BM-MSCs from healthy control groups with individual bacterial extract (BE) from each LBP+FR (n = 10) and LBP (n = 10) subject under various conditions for 48h. These BE were prepared from the least diverse fecal samples, based on α- diversity in LBP+FR and LBP groups. Then BM-MSCs from two groups were cultured in an adipogenesis medium for 14 days. Oil Red staining results revealed that BM-MSCs stimulated with BE from LBP+FR subjects promoted lipid droplet formation (Fig. 5A-B).

**Figure 5.**
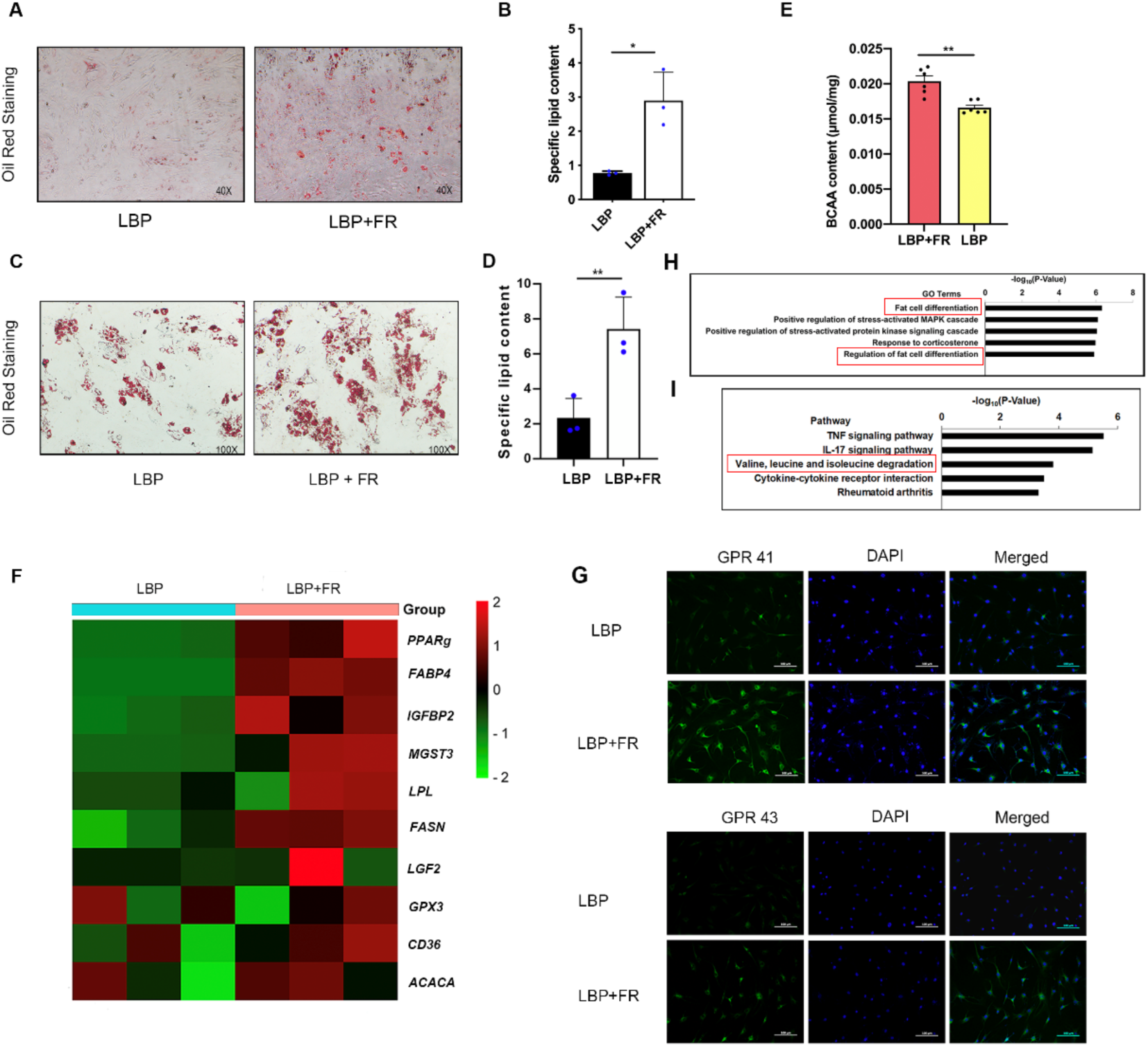
BM-MSCs from LBP subjects with FR exhibit a strong differentiation capability. (A-B) Representative images of Oil Red O staining of lipids (A) and quantification of the number of oil spots (B) in BM-MSCs under the bacterial extract’s stimulation from LBP+FR and LBP subjects. (C-D) Representative images of Oil Red O staining of lipids (C) and (D) quantification of the number of oil spots in BM-MSCs cultured in adipogenic induction medium for 14 days. 100× magnification. * indicates *p* <0.05; ** indicates *p* <0.01; and *** indicates *p* <0.001 by Wilcoxon rank-sum test. (E) The absolute qualification of BCAAs in BM-MSCs’ samples between LBP+FR (n = 6), LBP (n = 6) and HC groups (n = 6). (F) qRT-PCR analysis results of dysregulated adipogenic genes in LBP+FR and LBP derived BM-MSCs cultured in adipogenesis induction medium for 48 hours. (G) Immunocytochemistry of G protein-coupled receptor 41 (GPR41) and G-protein- coupled receptor 43 (GPR43) in BM-MSC. Scale bar = 100 μm. (H-I) GO and KEGG pathway enrichment analysis in LBP+FR (n = 6) and LBP groups (n = 6).

To examine the BM-MSCs’ differentiated potential in LBP+FR and LBP groups, BM-MSCs from these two groups were also cultured in an adipogenesis medium for 14 days. Oil Red staining showed that BM-MSCs from LBP+FR facilitated lipid droplet formation (Fig. 5C-D). We also measured the absolute qualification of BCAAs in BM-MSCs from LBP+FR and LBP groups. The level of BCAA in LBP+FR increased significantly compared with LBP groups. (Fig. 5E) Besides, RT-qPCR also confirmed the mRNA expression of adipogenic genes in BM-MSCs’ adipogenesis differentiation.

The expression of peroxisome proliferator–activated receptor-γ (Ppar-gama/PPARg) and fatty acid binding protein 4 (FABP4), 2 key markers of adipocyte differentiation, including other adipogenic differentiation markers (IGFBP2, MGST3, LPL and FASN) in BM-MSCs from LBP+FR patients, were higher than the BM-MSCs from LBP patients (Fig. 5F).

Besides, we found the fatty acids related to cell receptor G-protein-coupled receptor 41 (GPR41), also called free fatty acid receptor 3 (FFAR3) and G-protein-coupled receptor 43 (GPR43/FFAR2) is a Gαi -coupled receptor activated by SCFAs mainly produced from dietary complex carbohydrate fibers in the large intestine as products of fermentation by microbiota. This result showed that the BM-MSCs from LBP+FR patients had a stronger GPR41/43 immunostaining, which means BM-MSCs from LBP+FR patients were easily influenced by fatty acids. (Fig. 5G)

To further confirm the obtained results on the transcriptomic scale, whole transcriptome RNA sequencing was performed for the cells cultured in adipogenesis media solutions. Using gene ontology (GO) enrichment analysis, we observed that there were some enrichments in GO terms associated with fat cell differentiation and regulation of fat cell differentiation, this result illustrated that the BM-MSCs from LBP+FR patients have a promoted initiation of adipogenesis. (Fig. 5H) Furthermore, a KEGG pathway analysis in the upregulated gene group was also carried out. The top five related pathways of upregulated genes for the two groups are shown in Fig. 5I, including: TNF signaling pathway, IL-17 signaling pathway, Valine, leucine and isoleucine degradation, Cytokine-cytokine receptor interaction and Rheumatoid arthritis. It should be noted that the pathway about valine, leucine and isoleucine degradation was mentioned again, this pathway has been confirmed to have a close relationship with gut microbiome and serum metabolome in LBP+FR patients. Detailed results of GO and KEGG enrichment analysis are summarized in Supplementary Table 5.

These results show that BE from LBP+FR accelerates BM-MSCs’ adipogenesis, and BM-MSCs from LBP+FR group have stronger differentiated capability. These results kept consistent with the MRI pathology characteristics and indicated that BM-MSCs’ differentiation ability had a close relationship with the FR process. The BCAA degradation pathway was indicated to affect the development of FR positively.

### 7. BCAA degradation promotes BM-MSC’s adipogenic differentiation

We next detected the regulatory mechanisms of BCAAs on BM-MSC’s adipogenic differentiation. Genes related to BCAA degradation, including BCKDHA, BCKDHB, DBT, MCCC2, Pccb, Pcca and MUT were upregulated in BM-MSCs from LBP+FR. (Fig. 6A) As shown in Fig. 6B, the BCAA degradation pathway’s schematic illustration helps us know the roles of different enzymes involved in the BCAA pathway. Then, the changes of these key enzymes (BCKDHA, BCKDHB, DBT, MCCC2) were identified by immunoblot (Fig. 6C-D). Among them, the protein levels of BCKDHA complex, an important enzyme that controls the committed and initial steps of BCAA degradation to branched-chain acylcoA, was significantly upregulated in the LBP+FR group. Taken together, BCKDH’s activity was regarded as the most direct factor affecting the BCAA degradation in LBP+FR.

**Figure 6.**
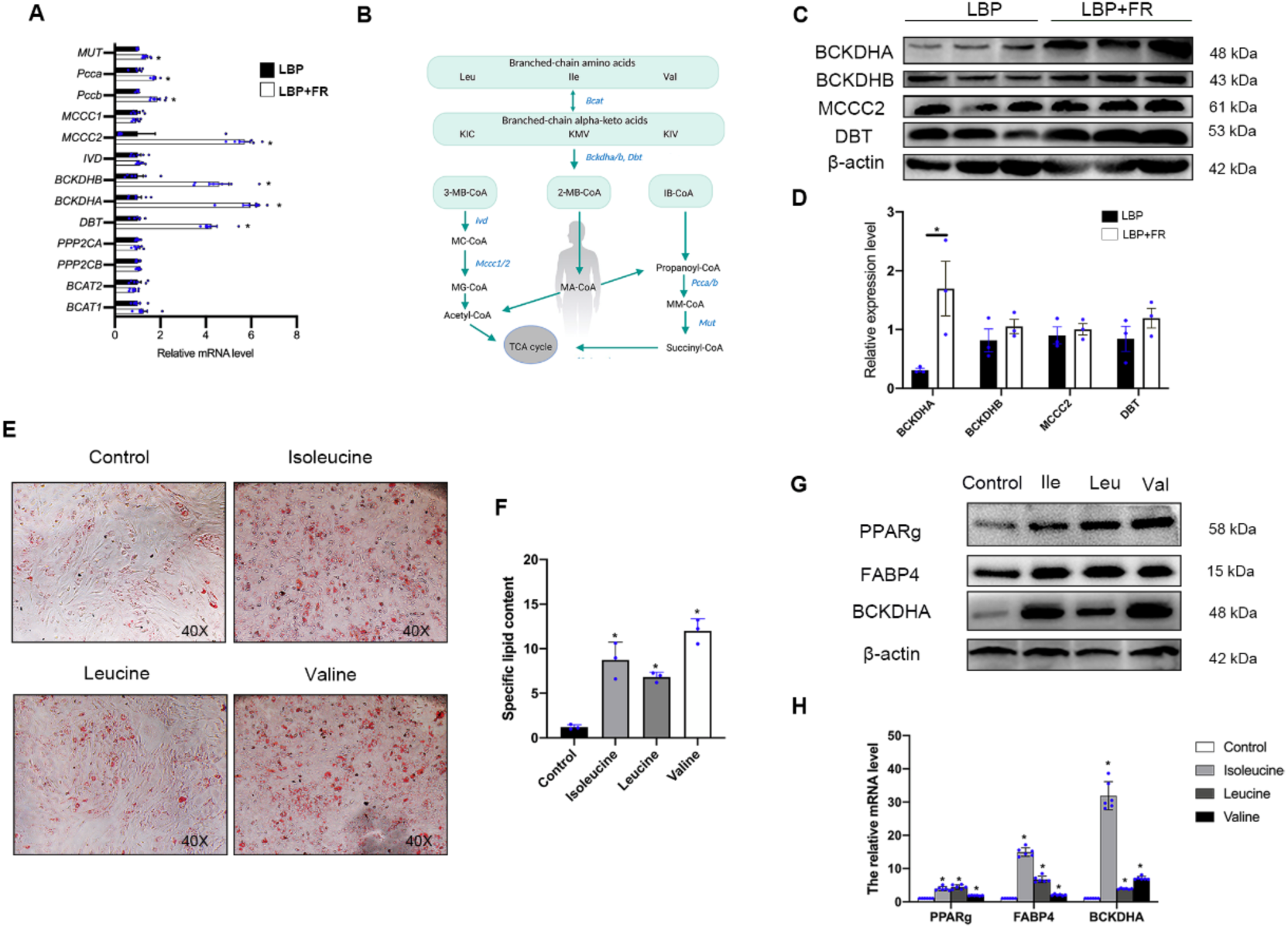
BCAA degradation positively regulates BM-MSC’s adipogenic differentiation. (A) RT-qPCR analyses of selected BCAA degradation genes in LBP+FR and LBP. (\**p* < 0.05, LBP+FR vs LBP, n =6). MUT, Methylmalonyl Coenzyme A Mutase; Pcca, Propionyl Coenzyme A Carboxylase, Alpha Polypeptide; Pccb, Propionyl Coenzyme A Carboxylase, Beta Polypeptide; MCCC1, Methylcrotonoyl-CoA Carboxylase 1; MCCC2, Methylcrotonoyl-CoA Carboxylase 2; IVD, Isovaleryl-CoA Dehydrogenase; BCKDHB, Branched Chain Keto Acid Dehydrogenase E1 Subunit Beta; BCKDHA, Branched Chain Keto Acid Dehydrogenase E1 Subunit Alpha; DBT, Dihydrolipoamide Branched-chain Transacylase E2; PPP2CA, Protein Phosphatase 2 Catalytic Subunit Alpha; PPP2CB, Protein Phosphatase 2 Catalytic Subunit Beta; BCAT2, Branched Chain Amino Acid Transaminase 2; BCAT1, Branched Chain Amino Acid Transaminase 1. (B) Schematic illustration of the BCAA degradation pathway. Enzymes examined in Fig. 6A are shown in blue. Degradation of Leu, Ile, and Val share the same initial steps catalyzed by Bcat2, Bckdha, Bckdhb and Dbt. Leu leucine, Ile isoleucine, Val valine, KIV α-ketoisovalerate, 3-MB-CoA 3-Methylbutanoyl-CoA, 2- MB-CoA 2-Methylbutanoyl-CoA, IB-CoA Isobutyryl-CoA, MC-CoA 2- Methylcrotonyl-CoA, MG-CoA 2-Methylglutaconyl-CoA, MA-CoA 2- Methylbutanoyl-CoA, MM-CoA Methylmalonyl-CoA. (C-D) Representative immunoblots of BCKDHA, BCKDHB, MCCC2, DBT and β- actin in BM-MSCs from LBP+FR and LBP groups (C) and statistical analyses of densitometric measurements of BCKDHA, BCKDHB, MCCC2 and DBT (D) are shown (\**p* < 0.05). (E-F) Representative images of Oil Red O staining of lipids (E) and quantification of the number of oil spots (F) in BM-MSCs from control and BCAA groups cultured in adipogenesis induction medium for 14 days. 40× magnification (G) Representative immunoblots of PPARg, FABP4, BCKDHA and β-actin in BM- MSCs from the BCAA group. (H) RT-qPCR analysis results of PPARg, FABP4 and BCKDHA in BCAA derived BM- MSCs cultured in adipogenesis induction medium for 14 days. (\**P* < 0.05).

To detect the correlations and functions of BCKDHA and BCAA in BM-MSCs’ differentiation, we tested the effect of the BCAA in an ex vivo cell culture model. We used three BCAAs (valine, isoleucine, leucine) to stimulate the BM-MSCs separately (Supplementary Fig. 6). BM-MSCs from BCAA treatment group exhibited increased adipogenesis (Fig. 6E-F), accompanied by increased protein levels and mRNA levels of PPARg and FABP4 (Fig. 6G-H). Moreover, BCKDHA protein and mRNA levels were significantly higher in the BCCA group compared with the control group. (Fig. 6G-H)

Notably, the untargeted metabolomics analysis of serum samples that show two metabolites (Thiamine pyrophosphate (Thpp) and 2-Methyl-1-hydroxypropyl-ThPP) related to the BCAAs degradation pathway were mainly enriched in the LBP+FR compared with the LBP. (Fig. 3 and Supplementary Table. 4) These two metabolites, especially the ThPP, could be required as a coenzyme for the E1 component of the BCKDH complex and can also activate the complex by inhibiting BCKDH kinase (BDK).^23^ Taken together, BCKDHA contributes significantly to the LBP+FR process. Thus, these observations strongly suggest that amino acid metabolism (mainly intracellular BCAA degradation) and the critical enzyme BCKDHA play an essential role in BM-MSC’s adipogenesis in LBP+FR.

### 8. SIRT4 boosts adipogenesis and the expression of BCKDHA

Next, we explored the upstream regulatory mechanisms mediating the BCAA degradation pathway. Sirtuins are NAD^+^-dependent enzymes conserved from bacteria to human^13^ and three sirtuins-SIRT3, SIRT4 and SIRT5 localize to the mitochondrial matrix, are critical regulators of mitochondrial metabolic enzymes^24, 25^. Furthermore, proteomics research on fibroblasts has found SIRT4 can alter mitochondrial activity by binding to BCAA catabolic enzymes BCAT^26^ and MCCC1^27, 28^. Given that BCAA degradation is a mitochondrial process, we hypothesized that sirtuins might increase BM-MSC adipogenesis by regulating BCAA degradation (Fig. 7A).

**Figure 7.**
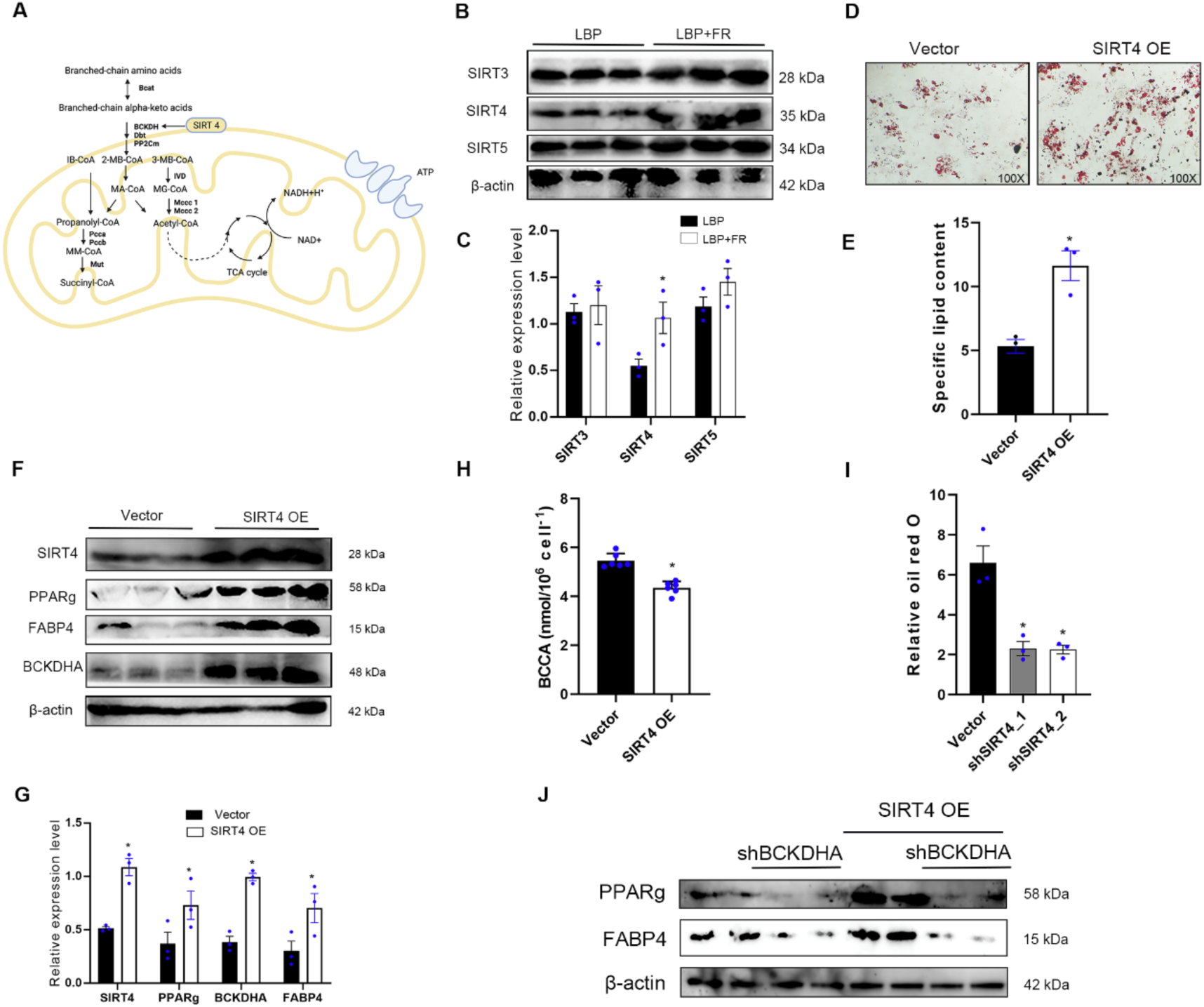
SIRT4 promotes adipogenesis and BCKDHA. (A) Model for SIRT4 regulation of BCAA catabolism. (B-C) Representative immunoblots of SIRT3, SIRT4, SIRT5 and β-actin in BM-MSCs from LBP+FR and LBP groups (B) and statistical analyses of densitometric measurements of SIRT3, SIRT4 and SIRT5 (C) are shown (\**p* < 0.05). (D-E) Representative images of Oil Red O staining of lipids (D) and quantification of the number of oil spots (E) in BM-MSCs from vector and SIRT4-overexpressing group cultured in adipogenesis induction medium for 14 days. 100× magnification (F-G) Representative immunoblots of SIRT4, PPARg, FABP4, BCKDHA and β-actin in BM-MSCs from vector and SIRT4-overexpressing groups (F) and statistical analyses of densitometric measurements of SIRT4, PPARg, FABP4 and BCKDHA (G) are shown (\**p* < 0.05). (H) The BCAA content in BM-MSCs from control and SIRT4-overexpressing groups after adipo-differentiation (\**p* < 0.05). (I) Quantification of relative oil red O in vector, shSIRT4_1 and shSIRT4_2 BM-MSCs differentiated for 14 days (\**p* < 0.05). (J) Western blot analysis of PPARg and FABP4 in BM-MSCs differentiated in14 days overexpressing SIRT4 or control vector and shBCKDHA or control plasmid.

After testifying by RT-PCR and WB, we found a largest increase in SIRT4 expression on protein and mRNA compared with the expression of SIRT3 and SIRT5 in BM-MSCs of LBP+FR patients (Supplementary Fig. 7A and Fig. 7B-C). To interrogate the role of mitochondrial SIRT4 in BM-MSCs’ differentiation, we overexpressed SIRT4 in normal BM-MSCs. SIRT4 overexpression caused a significant increase in adipogenesis, as assessed by oil red O staining (Fig. 7D-E). Besides, SIRT4 overexpression increased expression of genes and proteins related to adipogenesis (PPARg and FABP4) and BCKDHA (Supplementary Fig. 7B and Fig. 7F-G).

Given the established relationship between BCAA degradation and SIRT4, we hypothesized that SIRT4 might promote BM-MSC’s adipogenesis by regulating BCAA catabolism. We detected that SIRT4 overexpression decreased the BCAA content in BM-MSC after 14 days of adipo-differentiation (Fig. 7H). Moreover, we used two different shRNAs to knockdown SIRT4 (shSIRT4) and found that the loss of SIRT4 impaired adipocyte differentiation (Fig. 7I and Supplementary Fig. 7C). We also found that SIRT4 knockdown cells had significantly decreased protein and mRNA levels of BCKDHA compared to control BM-MSCs (Supplementary Fig. 7D and 7E). To further detect if BCKDHA is necessary for SIRT4-mediated adipogenesis in BM-MSCs, knocked down BCKDHA was used in the context of SIRT4 overexpression. We found that BCKDHA knockdown suppresses SIRT4-mediated increases of PPARg and FABP4 at protein level, suggesting that SIRT4 promotes adipogenesis through BCKDHA (Fig. 7J). Overall, these observations strongly suggest that SIRT4 positively regulates BCAA degradation and adipogenesis in BM-MSCs.

### 9. Gut microbiome- and metabolome-based prediction of LBP and FR

To determine whether differences in gut microbiome composition can be regarded as recognition biomarkers for distinguishing LBP patients from healthy participants, we trained Radom Forest (RF) classifiers on relative abundances of 16S tag sequences shared between the two LBP and HC groups. RF model was generated, and the corresponding receiver operating characteristic (ROC) curve was drawn to evaluate its distinguishing ability. As shown in Figures 8A and 8B, the classifiers yielded nearly perfect predictions (95% area under the ROC curve [AUC]) of LBP. This result shows that the random forest model based on fecal microbiota can distinguish LBP patients from healthy individuals, indicating that intestinal microbiota information can be used to identify LBP patients. In addition, a second RF classifier was trained to predict LBP with FR by the top 20 genus (16S genus) and the top 61 metabolites selected by all data constructed models. 10-fold cross-validation was applied to reduce the over-fitting effects. By combining the 16S top genera and the top 61 metabolites, the prediction power can achieve as high as an AUC of 0.94, demonstrating that microbial signatures and metabolites can accurately predict the FR in LBP patients. (Fig. 8C-E)

**Figure 8.**
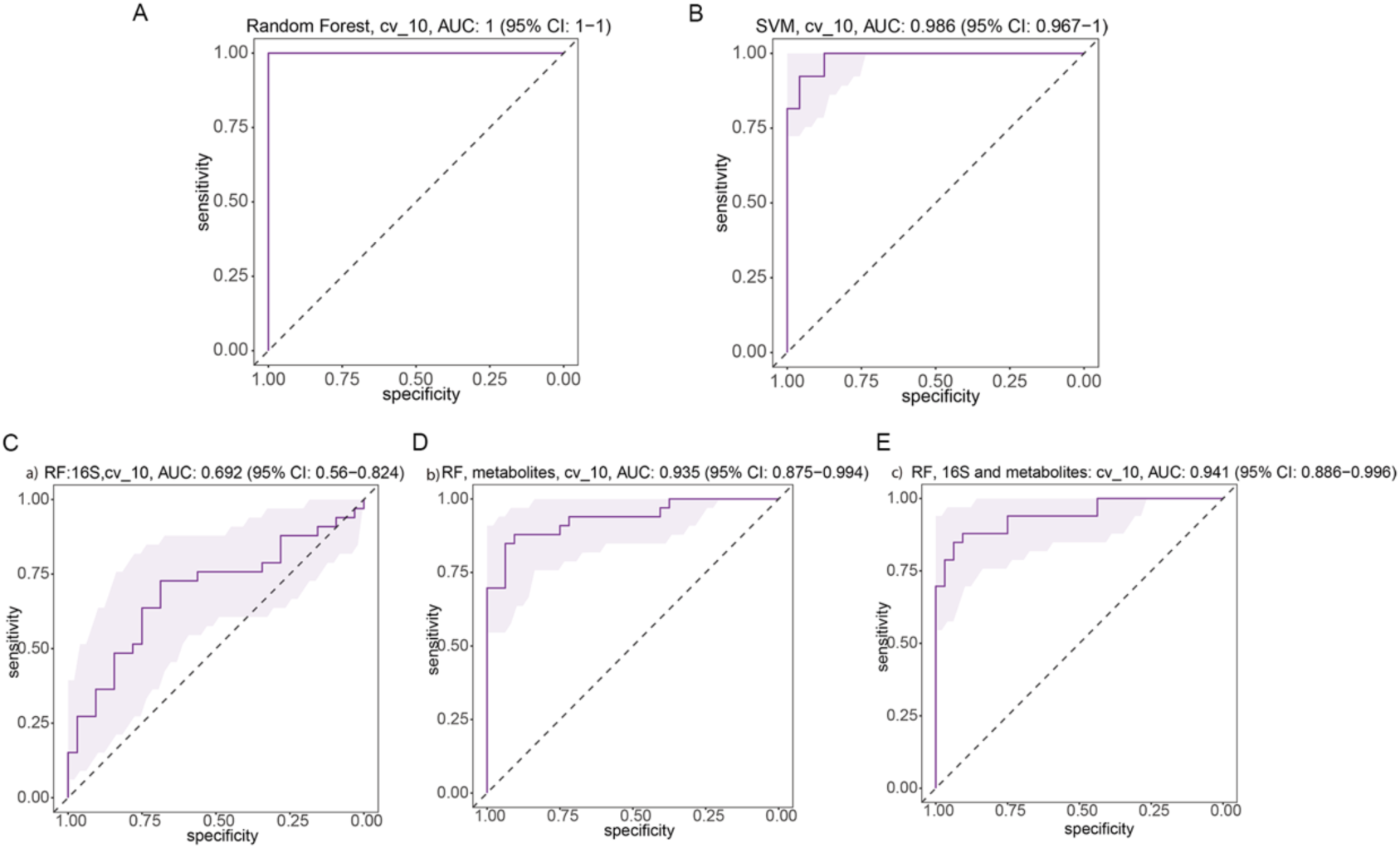
Random forest models to predict LBP and FR. (A-B) Prediction models to predict LBP to healthy control. The microbial signatures of 16S genera have AUC of 1 with 10-fold cross-validations for random forest, for support vector machine model, it has an AUC of 0.986. (C-E) Random Forest models to predict LBP types by top 20 16S genus and top 61 metabolites selected by all data constructed model.

We also found that RF model could discriminate LBP patients with FR from LBP subjects with an area under the curve ranging from 0.69 to 0.94 (all 16S genus, AUC = 0.69; all 10,000 metabolites, AUC = 0.935; combination of all 16S genus and metabolites, AUC = 0.94; Supplementary Fig. 8A-C).

The 16S has a very low AUC of around 0.6, but the metabolites can accurately predict types of LBP with an AUC of 0.88. While combining them together for all the data, the prediction power does not improve too much compared with metabolites only. These results demonstrated that these models based on gut microbiome and serum metabolites could distinguish LBP patients from healthy individuals and differentiated LBP patients with FR from LBP patients without FR in our cohort, suggesting that the gut microbiome and serum metabolite information could be applied to identify patients.

## Discussion

Researchers found chronic LBP with FR shares similar basic pathogenetic characteristics with osteoarthritic (OA), including MRI modalities, prevalence, pain, joint degeneration, risk factors, suggested etiologies and natural history with bone marrow lesions.^29^

There has been abundant evidence to support that gut microbiota, and its metabolites have been proposed as cofactors in OA^30^. However, few studies have directly examined gut dysbiosis’s effects on BMLs^13, 31^. Here, we outlined the landscapes and interaction networks of differential bacterial species and serum metabolites in the LBP with FR gut ecosystem. Disturbance of amino acid metabolism was a hallmark in the gut ecosystem of LBP with FR. Moreover, we utilised an ex vivo cell culture model to demonstrate an LBP+FR-specific effect of microbiota and metabolites on the BM- MSC’s adipogenesis differentiation, showing important clinical implications. Previous gut microbiome studies focused on the differential bacteria between LBP and non- LBP13 and did not clarify how the gut microbiome affects disease development. Our research compared the microbiome composition between LBP+FR, LBP and healthy controls using two methods:16S rRNA and metagenomics sequencing. 16S rRNA sequencing and metagenomic sequencing were prerequisites for screening the critical species associated with LBP with/without FR onset and identifying diagnostic markers for clinical applications.

16S rRNA sequencing results showed that the bacterial composition of LBP with/without FR differed from that in HC. LBP was characterized by enriched families Streptococcaceae, Prevotellaceae, Rikenellaceae, Alcaligenaceae, Clostridiaceae and Enterobacteriaceae; reduced families Ruminococcaceae and Lachnospiraceae. At the genus level, LBP with/without FR was distinguished from HC by the expansion of Prevotella, Escherichia, Megamonas and SMB53, and the reduction in Fecalibacterium, Roseburia, Lachnospira and Sutterella. Even though we did not find the different microbiome composition at family and genus level in the top abundant species between the LBP+FR group and LBP group, abundance comparisons of other not prominent genera showed that enriched Atopobium, Catenibacterium and Eubacterium and reduced Aggregatibacter in LBP+FR patients compared to LBP. Only a portion of the gut microbiota community revealed by shotgun sequencing could be detected by 16S rRNA gene sequencing.^32^ Shotgun sequencing, in particular, has more remarkable sensitive ability to identify taxa with less abundant than 16S rRNA gene sequencing when a sufficient number of reads are available. Furthermore, the alterations in metagenomes relate to disease features, the results involved could be utilized as therapeutic targets or the outputs used as fecal biomarkers, although this would need clinical and experimental validation. We demonstrated that the less abundant genera detected only by shotgun sequencing are biologically meaningful and can discriminate between three groups.

Our metagenomic analysis identified seven bacterial species linked with LBP (LBP with/without FR). Metagenomic microbiome sequencing data support an emerging core microbiome signature that LBP was characterized by reduced species of *Bilophila unclassified, Eubacterium hallii, Adlercreutzia equolifaciens* and *Lachnospiraceae bacterium 5 1 63FAA* compared to HC. *Adlercreutzia equolifaciens* has been proven to play a significant role in gut microbiota–host interactions, especially LBP^13^. *Adlercreutzia equolifaciens* is a well-known species that produce equol.^33^ In menopausal and postmenopausal women, equol has been proven critical in reducing bone loss and alleviating muscle and joint pain.^34^ Besides, Marloes et al. ^13^ used 16S rRNA analyses of fecal specimens in a group of 36 overweight or obese people who had or had not back pain. In contrast to our findings, Marloes et al. reported that the genera Adlercreutzia was found in greater abundance in people with back pain than in people who did not have back pain (*P* = 0.0008). These disparate findings are plausibly due to significant differences in the study cohorts. In Marloes’s study, all participants were overweight or obese people who additionally had a higher BMI (≥ 25 kg/m^2^), compared to our study cohort with a common BMI (≤ 24 kg/m^2^). In addition, other characteristics, including demographic, clinical, biochemical characteristics, diet, and physical activity, are different between Marloes’s study and our research.

Bilophila belongs to the Desulfovibrionaceae family, which produces hydrogen sulphide (H2S). Metabolises sulphated substances to create H2S, which can cause inflammation, cause epithelial cell genotoxicity and cytotoxicity, and disrupt intestinal barrier function^35^. Importantly, H2S has been reported to exert antinociceptive effects^36^ and sulfidogenic bacteria have recently been linked to the aetiology of chronic metabolic diseases ^37^. *Eubacterium hallii* and *Lachnospiraceae bacterium 5 1 63FAA*, with its ability to create large levels of SCFAs such as butyrate and propionate, have been hypothesised to be vital in maintaining gut microbial metabolic health and homoeostasis.^38, 39^ Although there is no clear evidence about the effects of SCFAs on LBP, SCFAs have been demonstrated to affect the inhibition of bone resorption or osteoclast formation either via activation of G Protein-Coupled Receptors (GPCR) or through histone deacetylase (HDAC) inhibition.^40^ Obviously, sufficiently powered studies in large cohorts are still needed to determine the specific alterations of the microbiome with LBP.

This paper focused on the differentiated species between LBP+FR and LBP. Three species, including identified enriched *Ruminococcus gnavus*, depleted *Roseburia hominis* and *Lachnospiraceae bacterium 8 1 57FAA* to distinguish LBP+FR from LBP and HC groups. *Ruminococcus gnavus* is the only enriched species in LBP+FR compared to LBP and HC groups. In most studies about *Ruminococcus gnavus*, an increased relative abundance of *Ruminococcus gnavus* was mainly linked with Crohn’s disease (CD), a major inflammatory bowel disease.^41^ It is worth noting that musculoskeletal pain occurs in 9–53% of IBD patients and is considered the most common extraintestinal manifestations.^42^ Although it is hard to connect the musculoskeletal pain and IBD through the current knowledge about *Ruminococcus gnavus*, these research findings provide the key to understanding the potential function of *Ruminococcus gnavus* in musculoskeletal diseases. Despite numerous studies associating *Ruminococcus gnavus* with various inflammatory diseases,^43, 44^ no molecular mechanisms to explain these correlations have been developed, and here we identified a possible mechanism for LBP+FR and *Ruminococcus gnavus*. Further proving that these critical species in the gut microbiome played a driving role in FR requires more *in vivo* experiments such as fecal microbiota transplant (FMT).

Gut bacteria are now well recognized as having a considerable influence on various metabolic pathways in the host. Both statistical correlation (co-expression network) and metabolic function support this conclusion in our study. In co-occurrence analysis among the gut microbiome and metabolites, changed microbiome species, *Ruminococcus gnavus* and *Roseburia hominis*, were substantially correlated with serum metabolites involved in most pathways (Fig. 4). Co-occurrence analysis showed the connection between BCAA degradation and *Ruminococcus gnavus*/*Lachnospiraceae bacterium 8 1 57FAA*. Besides, in metabolic analysis, the BCAAs degradation pathway was the most enriched pathway in LBP+FR group. This finding is consistent with the functional analysis of fecal microbiome based on metagenomics sequencing (Fig. 2I), KEGG pathway analysis showed BCAA pathway with functions related to metabolic pathways and signal transduction in LBP+FR group. Functional assignments and network analysis revealed that BCAA degradation pathway serves as the metabolic mediator for the cross-talk between the host and the microbiome. Thus, BCAA related pathways would differ in the disease condition compared to the healthy state. BCAA have key physiological roles in regulating protein synthesis, metabolism, food intake and ageing.^45^ BCAA also strongly correlates with metabolic diseases, including metabolic syndrome, type 2 diabetes, and urea cycle disorders.^45, 46^ LBP with FR could be characterised as a lipid metabolism disorder in the bone marrow. Even though we have demonstrated the relationship between BCAA and gut microbiome through co-occurrence analysis of the metagenomic sequencing and untargeted metabolome, more direct evidence is needed to establish their correlation.

Interestingly, Marloes et al. also found that overweight or obese individuals with back pain had a higher abundance of the genera Roseburia than those without back pain.^13^ On the one hand, this result confirms the prominent role of Roseburia in LBP. On the other hand, combined with the results of our study, the enrichment of Roseburia was changeable in different LBP conditions, like LBP+obesity or LBP+FR, which means Roseburia abundance could affect various metabolic pathways in LBP. Besides, Roseburia has been detected to associate with several diseases, including irritable bowel syndrome, ulcerative colitis, obesity, type 2 diabetes, nervous system disorders, and allergies. ^47, 48^ However, it must be acknowledged that these clinical findings are preliminary, and the impact of environmental and demographic factors cannot be ruled out.

Co-occurrence analysis found that the different microbial species and serum metabolites in LBP group were consistently mapped into Porphyrin and chlorophyll metabolism. Porphyrins are a class of metabolites that play a role in manufacturing life- sustaining chemicals^49^. The levels of bacterial porphyrin on the skin were shown to be substantially greater in acne vulgaris (acne) patients^50^. The dominant porphyrin species produced by *P. acnes*, coproporphyrin III, was shown to induce Staphylococcus aureus aggregation and biofilm formation in the nostrils^51^, which means *P. acnes* could be a preliminary cause of severe sepsis caused by Staphylococcus aureus. All information points to the connection between *P. acnes* and Porphyrin and chlorophyll metabolism. More interesting, researchers have found evidence that supported the close relationship between the LBP (especially the LBP+FR group) and *P. acnes*.^52^ In fact, there have always been doubts that the *P. acnes* in disc samples from LBP patients belong to sample contamination.^53^ Our results provide possible clues and future directions about serum metabolites clues in the relationship between LBP and *P. acnes*. However, skin microbial communities’ functions, metabolic activities, and host interactions are still not well understood.

The MRI signal intensity of the bone marrow in LBP with FR reflects a lipid conversion. BM-MSCs are stem cells and can differentiate into adipocytes or osteoblasts in the bone marrow.^54^ BM-MSCs tend to differentiate into adipocytes rather than osteoblasts with ageing, leading to progressive fat accumulation and bone loss.^55^ We demonstrated that BE from LBP+FR and LBP patients have different ability to induce BM-MSCs adipogenesis. And BM-MSCs from patients with LBP+FR exhibit a strong adipogenesis capability. Connected to the characteristics of MRI image and RNA sequencing results, BM-MSCs’ differentiated function plays a critical role in fatty replacement in bone marrow from LBP+FR patients. Moreover, we found that the expression of fatty-acids related cell receptor GPR 41/43 on BM-MSCs from LBP+FR patients was higher than the BM-MSCs from LBP patients. BM-MSCs from LBP+FR patients were easier to influence by some fatty acids produced by the gut microbiome. Notably, the differentiated microbiome in LBP+FR, *Ruminococcus gnavus* belongs to the family Ruminococcaceae, and *Roseburia hominis, Lachnospiraceae bacterium 8 1 57FAA*, belong to family Lachnospiraceae, are proved to be the main butyrate producing-bacteria in the human gut.^56^ These results shown gut microbiome in LBP+FR patients has the potential to regulate BM-MSCs adipogenesis through SCFAs. We did not find evidence of SCFAs from untargeted blood metabolism. To further detect the relationship between SCFAs and FR in LBP, targeted metabolomics about SCFAs should be done in the future.

Even though BM-MSCs’ differentiate in response to many factors, little is known about how metabolism drives differentiation. Here, we demonstrated that certain amino acids, particularly BCAAs, are consumed during BM-MSCs’ differentiation. BCAA catabolism supports mitochondrial respiration at this stage of BM-MSCs’ adipogenesis. The process of BM-MSCs differentiated into adipocytes in bone marrow was ATP- dependent. BCAA-induced mitochondrial respiration in adipogenesis may facilitate ATP generation for lipogenesis. These observations point to the importance of mitochondria in BM-MSCs’ differentiation process. Our study highlights the fundamental role of mitochondria as the site of providing enough energy to BM-MSCs’ adipogenesis.

Importantly, SIRT4 and induction of BCAA catabolism amplify the expression of PPARg and FABP4 and its targets, including BCAA catabolism enzymes: BCKDHA. Although, it is unclear how SIRT4 function in the mitochondria impacts transcriptional activity of PPARg and FABP4. The work from Laurent et al. showed that the loss of SIRT4 increases expression of PPARa target genes in fatty acid catabolism^57^. Mechanistically, in the absence of SIRT4, NAD+ levels are elevated and activate SIRT1, which can then be recruited to PPARa and co-regulate transcription. Previous work identified the link between SIRT4 and BCAA catabolism through MCCC1 in the liver^58^. Here, we find that SIRT4 enhances BCKDHA activity in BM-MSCS and that loss of this regulation results in decreased adipogenesis. SIRT4 has been reported to bind to numerous substrates involved in BCAA catabolism, including MCCC1, BCKDHA, DBT and dihydrolipoyl dehydrogenase (DLD)^28, 58^, raising the possibility that SIRT4 engages this pathway at multiple enzymatic nodes. Additional investigation is required to determine which SIRT4-BCAA catabolism enzyme interactions are most important for promoting adipogenesis in BM-MSCs from LBP+FR patients. Given that numerous enzymatic activities of SIRT4 have been revealed^59^, SIRT4 may augment BCKDHA activity in BM-MSCs through a previously unidentified activity.

In summary, using multi omics data, we have presented evidence that LBP or LBP+FR were characterized by gut bacterial disturbances and serum metabolites, representing the overall disturbances of LBP or LBP+FR gut ecology. Furthermore, disturbance of microbial amino acid metabolism (BCAA degradation pathway) was a potential source of disease biomarkers in LBP+FR. Moreover, this work identifies a significant process in adipogenesis regulated by SIRT4-mediated stimulation of BCAA metabolism with important developmental implications in BM-MSCs. These findings provide new directions to uncover pathogenesis and develop novel diagnostic strategies for chronic LBP with FR.

## Material and methods

### Patients’ samples

We recruited 107 chronic LBP Patients with/without FR and 31 healthy controls. The ethics approval was obtained from the Human Research and Ethics Committee of the First Affiliated Hospital (no: 2021-270), Sun Yat-sen University, China. For the chronic LBP group, participants were recruited for this project if they were any gender aged between 20-70 years, willing to undergo an MRI of the lumbar spine for diagnosing FR, had chronic LBP at least half the day in the past six months, and no previous low back surgery, and had not been diagnosed with ankylosing spondylitis, osteoporotic vertebral fracture, multiple myeloma, metastatic carcinoma, and psoas abscess. Healthy individuals were adults with no history of LBP and FR or spinal disease. We placed the advertisement for participant recruitment (participants with LBP or NO LBP) at the First Affiliated Hospital. Participants were excluded from the project if they had any antibiotics or oral corticosteroids in the past month; known gastrointestinal disease; previous gastrointestinal surgery; regular proton-pump inhibitor, or lactulose therapy. These drugs and therapies will interfere with the gut microbiome composition.

### Specimen collection and library preparation

All participants were given a fecal collection kit and guided to generate their faces within two days. All fecal samples were frozen and stored at -80℃. Microbial DNA was extracted from stool by MOBIO PowerSoil^®^ DNA Isolation kit (MOBIO Laboratories, Carlsbad, CA, USA) with bases-beating step following the manufacturer’s protocol. Sequencing libraries were generated using NEB Next^®^ Ultra™ DNA Library Prep Kit for Illumina^®^ (New England Biolabs, MA, USA) following the manufacturer’s recommendations and index codes were added. The library was sequenced on an Illumina NovaSeq 6000 platform and 150 bp paired end reads were generated.

### 16S rRNA gene sequencing analysis

We merged, applied quality control and clustered the 16S rRNA gene reads into operational taxonomic units (OTUs) using QIIME 2 Taxonomic groups were based on the Greengenes Database V.13_8 using closed reference to perform referenced-based OTU clustering.^60, 61^ Values for alpha diversity (Chao1 Index, Shannon’s Index, Phylogenetic Diversity (PD) Whole Tree Index and observed OTUs), beta diversity (Unweighted UniFrac distance metrics) and principal coordinate analysis (PCoA) employed based on the Bray_Curtis distance, Unweighted_Unifrac and Weighted_Unifrac distance were generated by QIIME 2 Permutational multivariate analysis of variance was performed to determine if the compositions of microbiota differed between groups. Linear discriminant analysis effect size (LEfSe) was performed to determine the features most likely to explain the differences between groups.

### Metagenomic sequencing analysis

The raw data processing using Trimmomatic^62^ (v.0.36) was conducted to acquire the Clean Data for subsequent analysis. MEGAHIT (Version v1.0.6) was used to assemble the metagenomics Clean Data. Scaftigs were obtained by mixing assembled scaffolds from N connection. The contigs with a length of or more than 500 bp were selected as the final assembling result. Open reading frames were predicted from both single and mixed Scaftigs using MetaGeneMark (Version 3.38) and filtered the length information shorter than 90 nt from the predicted result with default parameters. Then CD HIT (Version:4.7) is adopted to remove redundancy and obtain the unique initial gene catalogue. All predicted genes with a 95% sequence identity and 90% coverage were clustered. Reads and number of reads in each sample were obtained by using BBMAP software.

We annotated gene sets using DIAMOND software based on the NCBI NR database. Each gene is assigned to the highest-scoring taxonomy based on a unified database. This achieves simultaneous assessment of the gut microbiome of LBP patients with or without FR. The DIAMOND software is adopted to blast Unigenes to KEGG annotation. Alpha-diversity analysis, including Chao-1-richness index, Observed-otus, Shannon’s diversity index and PD-whole-tree index was utilized conducted and visualized using the fossil and vegan packages in R. PCoA was used to visually evaluate the overall difference and similarity of gut bacterial communities between the LBP patients with or without FR and HC groups.

Group differences were tested by using the PERMANOVA. The differential bacterial species between the three groups were identified using LEfSe with linear discriminant analysis (LDA) score >2. Furthermore, Wilcoxon rank-sum test was used to identify the differential species and metabolites between different groups (false discovery rate, <0.05). Then, the correlated microbial genes in the KEGG pathway were detected.

### Serum and BM-MSCs preparation

Serum samples were collected from the whole blood by serum-separator tubes, and serum samples should be stored at -80℃ within 2 hours. BM-MSCs samples were immediately separated from the bone marrow and stored at -80℃. Sample treatments followed the recommended processing guides for untargeted metabolomic study.

### Metabolomics measured by LC-MC

Untargeted metabolomics Profiles were performed by the Beijing Genomics institution (BGI). Advanced mass spectrometer Xevo G2-XS QTOF (Waters, UK) was used to compare the three groups’ serum metabolites signatures. Commercial software Progenesis QI (version 2.2) (Waters, UK) and the BGI’s metabolomics software package metaX were used for mass spectrometry data analysis, wherein identification was based on the KEGG database. PCoA was applied to discriminate the samples from different groups visually. The cluster analysis was performed to group the selected differential metabolites. Metabolic pathway in which the differential metabolites involved were enriched referred to the KEGG pathway and database.

### Metabolomic and metagenomic data integration

Correlation analysis between metagenomic and metabolomic data was undertaken using genomes and metabolites identified as significantly different between LBP+FR, LBP and healthy samples incorporating adjustment for age, sex and BMI. The co-occurrence among the gut microbiome and serum metabolites was calculated based on the relative abundances by Spearman’s rank correlation coefficient (*P* < 0.05). The network layout was calculated and visualized using a circular layout by the Cytoscape software. Only edges with correlations greater than 0.5 were shown in the bacteria and serum metabolites, and unconnected nodes were omitted. Correlation coefficients with a magnitude of 0.5 or above were selected for visualization in Cytoscape.

### Branched chain amino acid assay

Following the manufacturer’s instructions, a commercial colorimetric measurement kit (ab83374, Abcam, United Kingdom) was utilized to quantify the BCAA levels in serum and cell culture media samples. The serum samples were diluted 2.5 times and incubated at room temperature for 30 minutes. The absorption at 450 nm was measured with a microplate reader, and then the BCAA’s level was calculated using the standard curve.

### BM-MSCs-isolation and ex vivo culture

The mononuclear cells (MNCs) were extracted from the marrow cavity of the intervertebral body according to a widely used protocol.^63^ Briefly, the collected bone marrow was gently mixed with PBS (1:2). The mixture was equally layered onto an equal volume of Ficoll (1,077 g/m, GE health care, Chicago, USA) and collected buffy coat by using a density gradient centrifugation. Then the isolated buffy coat was washed twice with PBS and spun with centrifugation. The gotten red blood cells were seeded in DMEM low glucose with 10% FBS (Gibco, Life Technologies Ltd, Paisley, UK) at a density of 160,000/cm^2^. All non-adherent cells were removed. The culture medium was changed twice a week, and the BM-MSCs could be harvested by the trypsin (Thermofisher, USA) after about 3∼5 days and transferred to a new flask.

### Bacterial extract preparation for stimulation of BM-MSCs

50 mg of fresh fecal was suspended in 1.5 mL of PBS and filtered through a 40 μm cell strainer to get bacterial extracts (BE). Fecal samples were obtained from subjects with LBP+FR and LBP controls. After that, samples were reconstituted in 1.5 mL of PBS containing protease inhibitor (Roche) and phosphatase inhibitor (Roche) supplements. After 10 minutes, samples were heat-inactivated at 65 °C for 1 hour after sonication. According to the manufacturer’s instructions, the protein concentration in the resulting suspension was determined using the Pierce BCA protein assay kit (Thermo Scientific), and the LPS levels were determined using the Pierce Chromogenic Endotoxin Quantification Kit (ThermoFisher).

Prior to BM-MSCs’ stimulation, all BE were prepared to protein concentration of 10 mg/mL, as higher concentrations were shown in dose-titration experiments to compromise BM-MSCs viability to <90%. Each subject’s individual BE was used to stimulate each BM-MSCs sample (i.e. self and non-self BE stimulation).

### Immune-Phenotype

BM-MSCs were counted and divided 1x10^5^/ tube and then re-suspended in 100 μL of antibodies mix. Subsequently, cells were incubated for 30 minutes at 4 degrees, washed and analysed with a FACSCanto flow cytometer (BD PharMingen) and with the FACSDiva software (Tree Star, Inc. Ashland, OR). BM-MSCs have characterized with some special monoclonal antibodies again CD105, CD29, CD34 and CD45 (Cyagen, #HUXMX-09011) associated with different fluorochromes.

### Oil Red O staining and quantification

BM-MSCs were cultured in the adipogenesis induction medium (α-MEM containing 0.5 mM 3-isobutyl-1-methylxanthine, 1 μM dexamethasone, 5 μg/ml insulin, and 10% FCS) for 14 days. The medium was replaced twice a week. Oil Red O was performed to detect the mature adipocytes. Briefly, BM-MSCs were washed twice in PBS, fixed with 3.7% formalin, washed with 60% isopropanol, and stained with 0.3% Oil Red O solution. The excess stain was removed by washing the cells with sterile water, and cells were then dried for imaging. For quantification, lipid droplets were solubilized in 100% isopropanol and quantified by determining the resultant absorbance at 492nm.

Cells were counterstained with 0.01% crystal violet and extracted with 100% methanol, and absorbance was measured at 570 nm.

### Gene expression analysis by RNA sequencing

Total RNA extraction was done from BM-MSCs after different treatments with TRIzol reagent. Subsequently, the RNA samples were sent to BGI Co., LTD (Shenzhen, China), and BGISEQ-500 platform was applied to perform RNA-sequencing. Differentially expressed genes were determined based on Q value (Adjusted P-value) <= 0.05.

### qRT-PCR analysis

Total RNA from BM-MSCs was extracted by the TRIzol reagent (Invitrogen). The RNA concentration was measured by the Nanodrop 2100, and reverse transcription was performed using 1 μg total RNA. Total RNA was reverse-transcribed into the first-strand cDNA using the Superscript First-Strand Synthesis Kit (Invitrogen). cDNA transcripts were quantified by Rotor Gene Real-Time PCR System (Qiagen) using SYBR Green (Biorad). Primer sequences were shown in Table 2.

**Table 2.**
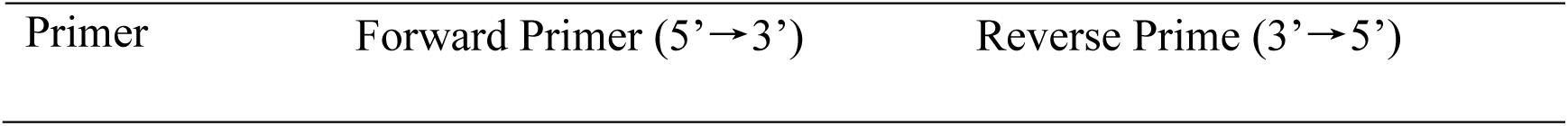
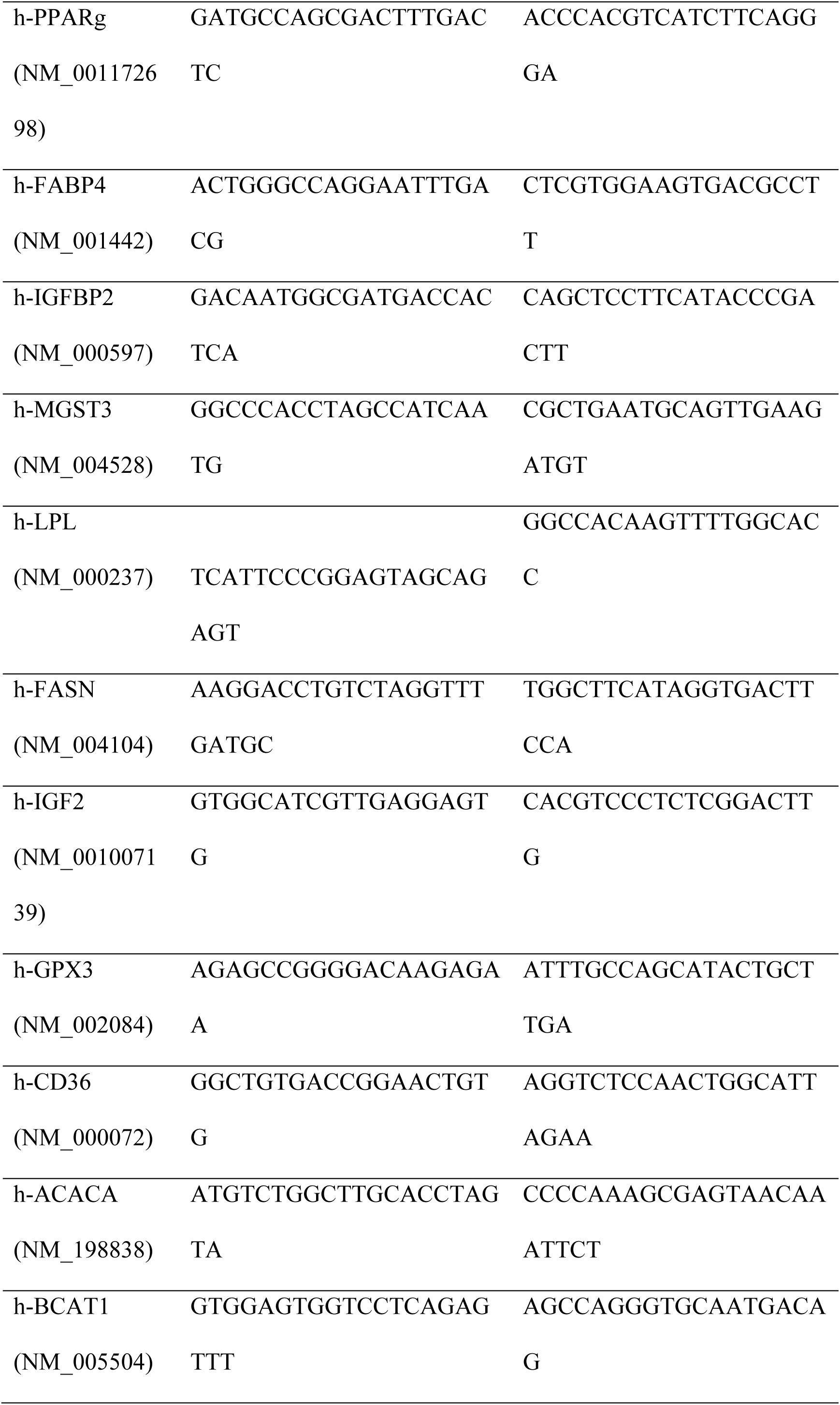
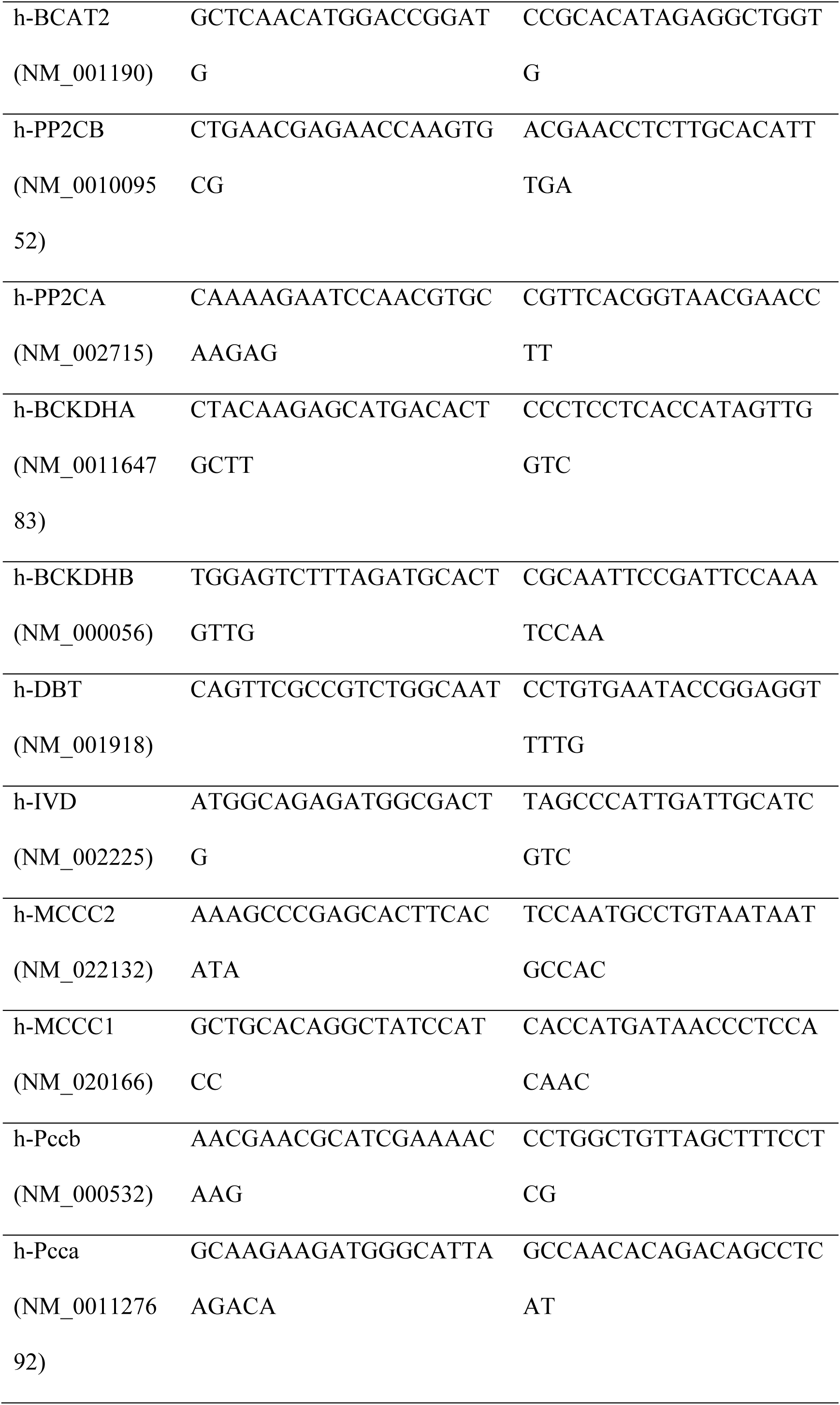
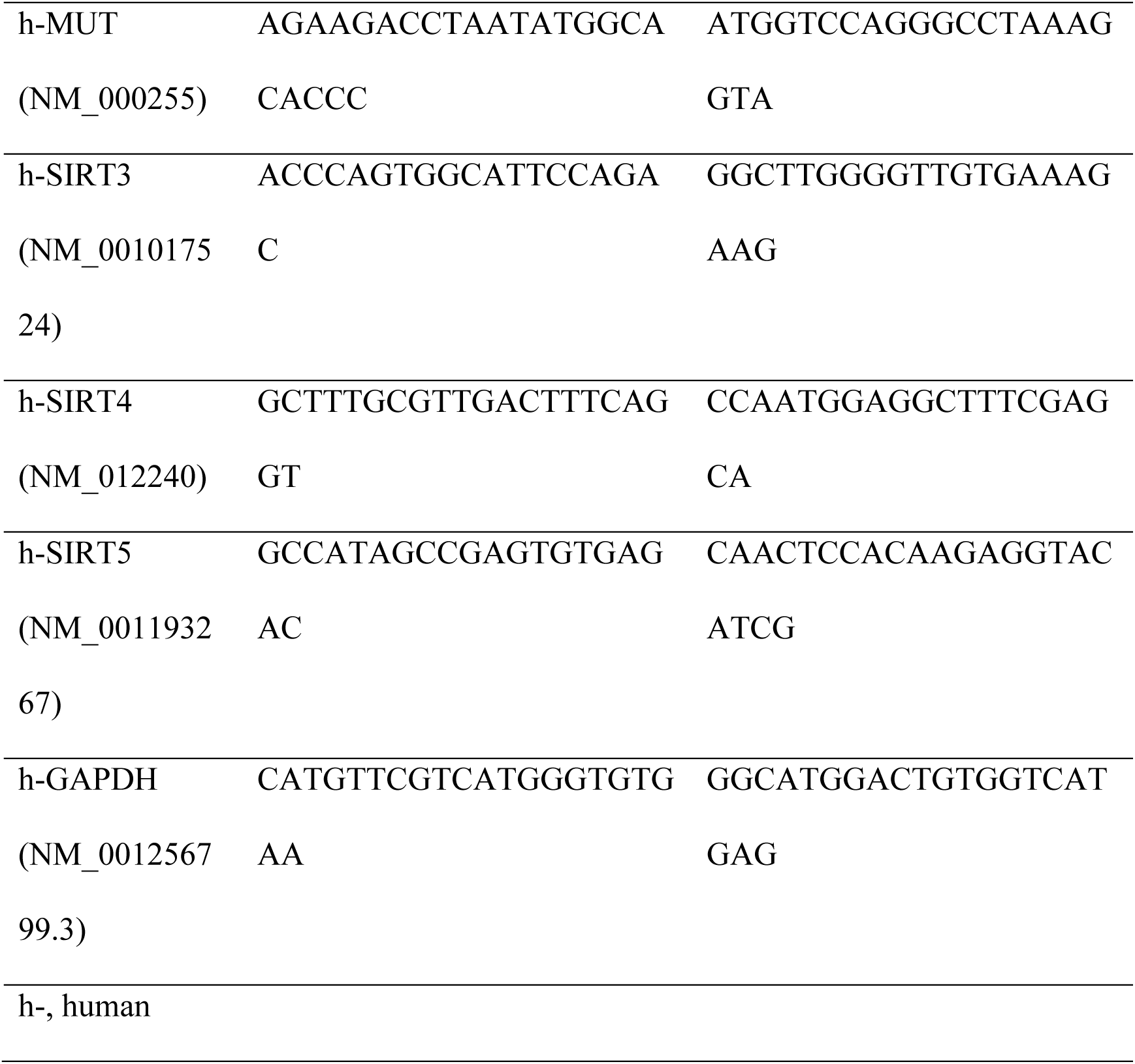
The Primer Sequences Used for Quantitative Real-Time PCR.

### BM-MSC Transfection

BM-MSCs were transfected by short hairpin RNAs (shRNAs). For overexpression studies, BM-MSCs’ overexpressing SIRT4 were established by the Lentiviral Packaging Kits (TR30037; ORIGEN) according to the manufacturer’s protocol. Stable overexpression of SIRT4 was achieved by infecting BM-MSCs with lentiviruses for 72 h followed by culturing in medium with 2 µg/ml puromycin (Beyotime Institute of Biotechnology, Haimen, China) at 37 °C and 5% CO2 for 2 weeks.

### Immunocytochemistry

BM-MSCs were washed with phosphate-buffered saline (PBS) three times, fixed with 4% paraformaldehyde for 15 min, permeabilized in 0.3% Triton X-100 for 10 min and blocked with 10% normal goat serum for 1 hour at room temperature. The following antibodies were used as primary antibodies: FFAR 3 (1: 100, #PA5-97745, Invitrogen) and GPR 43(1: 200, #BS-13536R, Thermo Scientific). Alexa Fluor 488 Dye-conjugated secondary antibody (Invitrogen) was used for detecting indirect fluorescence and then mounted on glass slides with Vectashield (Vector laboratories).

### SDS Page Immunoblot

BM-MSC lysates were prepared by RIPA lysis buffer (catalogue #P0013B; Beyotime Biotechnology, Shanghai, China) supplemented with PMSF protease inhibitor on ice for 30 min. Bicinchoninic acid (BCA) protein assay kit (catalogue #P0010S; Beyotime Biotechnology) was used to quantify the protein concentration according to the instruction. Cell proteins were isolated by the SDS-PAGE blotted on PVDF membranes (Millipore). And then blocked with 5% fat-free dry ilk at RT for 1 h. The PVDF membranes were incubated with specific antibodies to related genes at 4℃. Next incubated with appropriate HRP-conjugated secondary antibodies and exposed to x-ray films.

### Statistical analysis

The number of independent experiments can be found in the relevant figure legends. One-way analysis of variance (ANOVA) was used to compare continuous variables, which were displayed as means ± SD. Data of metagenome and metabolome were presented as means ± standard error of the mean (SEM). GraphPad Prism and excel were used for statistical analysis. The statistical significance level was set at a *P* value of < 0.05.

## Data availability

The 16S rRNA amplicon and metagenomic sequencing data have been deposited in National Center for Biotechnology Information (NCBI) with the primary accession code: PRJNA822996. The mass spectrometry raw data have been deposited on the MetaboLights (ID: MTBLS4644). The remaining data are available within the article, supplementary information or available from the authors upon request.

## Acknowledgements

We appreciate the department of Spine Surgery (The First Affiliated Hospital, Sun Yat-Sen University, Guangzhou) and the Department of Orthopedic Surgery (The First Affiliated Hospital of Soochow University, Suzhou, Jiangsu) for providing funding support.

We thank Dr. Kyle Sheldrick for his guidance and kind help to Wentian about human ethics application. SpineLabs is supported by in-kind support from St George Private Hospital and unrestricted donations from Nuvasive Australia and Baxter Australasia. Spine Service receives education and training grants from Globus Medical.

## Author Contributions

W.T.L conceptualized the project and planned, executed, and prepared the work for publication. ADD & AD initiated the study and along with Z.Z.M provided supports, supervision. W.T.L., J.T., J.J.Z., Q.Y., W.Y.D., S.D.Y., C.Y. and J.Z. collected samples, JZ’s lab and X.T.J provided analytical support. W.T.L and J.T. performed all experiments and produced all figures and tables. X.P.B. and K.T.L. helped to analyze the data.

## References

1. Knezevic, N.N., Candido, K.D., Vlaeyen, J.W.S., Van Zundert, J., and Cohen, S.P. (2021). Low back pain. Lancet 398, 78–92. 10.1016/s0140-6736(21)00733-9.

2. Thompson, K.J., Dagher, A.P., Eckel, T.S., Clark, M., and Reinig, J.W. (2009). Modic changes on MR images as studied with provocative diskography: clinical relevance--a retrospective study of 2457 disks. Radiology 250, 849–855. 10.1148/radiol.2503080474.

3. Jensen, T.S., Karppinen, J., Sorensen, J.S., Niinimäki, J., and Leboeuf-Yde, C. (2008). Vertebral endplate signal changes (Modic change): a systematic literature review of prevalence and association with non-specific low back pain. Eur Spine J 17, 1407–1422. 10.1007/s00586-008-0770-2.

4. Kuisma, M., Karppinen, J., Niinimäki, J., Ojala, R., Haapea, M., Heliövaara, M., Korpelainen, R., Taimela, S., Natri, A., and Tervonen, O. (2007). Modic changes in endplates of lumbar vertebral bodies: prevalence and association with low back and sciatic pain among middle-aged male workers. Spine (Phila Pa 1976) 32, 1116–1122. 10.1097/01.brs.0000261561.12944.ff.

5. Kuisma, M., Karppinen, J., Niinimäki, J., Kurunlahti, M., Haapea, M., Vanharanta, H., and Tervonen, O. (2006). A three-year follow-up of lumbar spine endplate (Modic) changes. Spine (Phila Pa 1976) 31, 1714–1718. 10.1097/01.brs.0000224167.18483.14.

6. Chen, Q., Shou, P., Zheng, C., Jiang, M., Cao, G., Yang, Q., Cao, J., Xie, N., Velletri, T., Zhang, X., et al. (2016). Fate decision of mesenchymal stem cells: adipocytes or osteoblasts? Cell Death Differ 23, 1128–1139. 10.1038/cdd.2015.168.

7. Dudli, S., Fields, A.J., Samartzis, D., Karppinen, J., and Lotz, J.C. (2016). Pathobiology of Modic changes. Eur Spine J 25, 3723–3734. 10.1007/s00586-016-4459-7.

8. van Gastel, N., and Carmeliet, G. (2021). Metabolic regulation of skeletal cell fate and function in physiology and disease. Nature Metabolism 3, 11–20. 10.1038/s42255-020-00321-3.

9. Tajik, N., Frech, M., Schulz, O., Schälter, F., Lucas, S., Azizov, V., Dürholz, K., Steffen, F., Omata, Y., Rings, A., et al. (2020). Targeting zonulin and intestinal epithelial barrier function to prevent onset of arthritis. Nat Commun 11, 1995. 10.1038/s41467-020-15831-7.

10. Li, J., Raizada, M.K., and Richards, E.M. (2021). Gut-brain-bone marrow axis in hypertension. Curr Opin Nephrol Hypertens 30, 159–165. 10.1097/mnh.0000000000000678.

11. Longstreth, G.F., and Yao, J.F. (2004). Irritable bowel syndrome and surgery: a multivariable analysis. Gastroenterology 126, 1665–1673. 10.1053/j.gastro.2004.02.020.

12. Whorwell, P.J. (2004). Back pain and irritable bowel syndrome. Gastroenterology 127, 1648–1649. 10.1053/j.gastro.2004.09.071.

13. Dekker Nitert, M., Mousa, A., Barrett, H.L., Naderpoor, N., and de Courten, B. (2020). Altered Gut Microbiota Composition Is Associated With Back Pain in Overweight and Obese Individuals. Front Endocrinol (Lausanne) 11, 605. 10.3389/fendo.2020.00605.

14. Urquhart, D.M., Zheng, Y., Cheng, A.C., Rosenfeld, J.V., Chan, P., Liew, S., Hussain, S.M., and Cicuttini, F.M. (2015). Could low grade bacterial infection contribute to low back pain? A systematic review. BMC Med 13, 13. 10.1186/s12916-015-0267-x.

15. Li, B., Dong, Z., Wu, Y., Zeng, J., Zheng, Q., Xiao, B., Cai, X., and Xiao, Z. (2016). Association Between Lumbar Disc Degeneration and Propionibacterium acnes Infection: Clinical Research and Preliminary Exploration of Animal Experiment. Spine (Phila Pa 1976) 41, E764–e769. 10.1097/brs.0000000000001383.

16. Li, W., Lai, K., Chopra, N., Zheng, Z., Das, A., and Diwan, A.D. (2022). Gut-disc axis: A cause of intervertebral disc degeneration and low back pain? European Spine Journal 31, 917–925. 10.1007/s00586-022-07152-8.

17. Dodd, D., Spitzer, M.H., Van Treuren, W., Merrill, B.D., Hryckowian, A.J., Higginbottom, S.K., Le, A., Cowan, T.M., Nolan, G.P., Fischbach, M.A., and Sonnenburg, J.L. (2017). A gut bacterial pathway metabolizes aromatic amino acids into nine circulating metabolites. Nature 551, 648–652. 10.1038/nature24661.

18. Wang, T.J., Larson, M.G., Vasan, R.S., Cheng, S., Rhee, E.P., McCabe, E., Lewis, G.D., Fox, C.S., Jacques, P.F., Fernandez, C., et al. (2011). Metabolite profiles and the risk of developing diabetes. Nat Med 17, 448–453. 10.1038/nm.2307.

19. Green, C.R., Wallace, M., Divakaruni, A.S., Phillips, S.A., Murphy, A.N., Ciaraldi, T.P., and Metallo, C.M. (2016). Branched-chain amino acid catabolism fuels adipocyte differentiation and lipogenesis. Nature chemical biology 12, 15–21. 10.1038/nchembio.1961.

20. Quince, C., Walker, A.W., Simpson, J.T., Loman, N.J., and Segata, N. (2017). Shotgun metagenomics, from sampling to analysis. Nature Biotechnology 35, 833–844. 10.1038/nbt.3935.

21. Tickle, T.L., Segata, N., Waldron, L., Weingart, U., and Huttenhower, C. (2013). Two-stage microbial community experimental design. The ISME Journal 7, 2330–2339. 10.1038/ismej.2013.139.

22. Clos-Garcia, M., Andrés-Marin, N., Fernández-Eulate, G., Abecia, L., Lavín, J.L., van Liempd, S., Cabrera, D., Royo, F., Valero, A., Errazquin, N., et al. (2019). Gut microbiome and serum metabolome analyses identify molecular biomarkers and altered glutamate metabolism in fibromyalgia. EBioMedicine 46, 499–511. 10.1016/j.ebiom.2019.07.031.

23. Noguchi, S., Kondo, Y., Ito, R., Katayama, T., Kazama, S., Kadota, Y., Kitaura, Y., Harris, R.A., and Shimomura, Y. (2018). Ca(2+)-dependent inhibition of branched-chain α- ketoacid dehydrogenase kinase by thiamine pyrophosphate. Biochem Biophys Res Commun 504, 916–920. 10.1016/j.bbrc.2018.09.038.

24. Vassilopoulos, A., Fritz, K.S., Petersen, D.R., and Gius, D. (2011). The human sirtuin family: evolutionary divergences and functions. Hum Genomics 5, 485–496. 10.1186/1479-7364-5-5-485.

25. Giblin, W., Skinner, M.E., and Lombard, D.B. (2014). Sirtuins: guardians of mammalian healthspan. Trends Genet 30, 271–286. 10.1016/j.tig.2014.04.007.

26. Lei, M.Z., Li, X.X., Zhang, Y., Li, J.T., Zhang, F., Wang, Y.P., Yin, M., Qu, J., and Lei, Q.Y. (2020). Acetylation promotes BCAT2 degradation to suppress BCAA catabolism and pancreatic cancer growth. Signal Transduct Target Ther 5, 70. 10.1038/s41392-020-0168-0.

27. Wirth, M., Karaca, S., Wenzel, D., Ho, L., Tishkoff, D., Lombard, D.B., Verdin, E., Urlaub, H., Jedrusik-Bode, M., and Fischle, W. (2013). Mitochondrial SIRT4-type proteins in Caenorhabditis elegans and mammals interact with pyruvate carboxylase and other acetylated biotin-dependent carboxylases. Mitochondrion 13, 705–720. 10.1016/j.mito.2013.02.002.

28. Mathias, R.A., Greco, T.M., Oberstein, A., Budayeva, H.G., Chakrabarti, R., Rowland, E.A., Kang, Y., Shenk, T., and Cristea, I.M. (2014). Sirtuin 4 is a lipoamidase regulating pyruvate dehydrogenase complex activity. Cell 159, 1615–1625. 10.1016/j.cell.2014.11.046.

29. Felson, D.T., Niu, J., Guermazi, A., Roemer, F., Aliabadi, P., Clancy, M., Torner, J., Lewis, C.E., and Nevitt, M.C. (2007). Correlation of the development of knee pain with enlarging bone marrow lesions on magnetic resonance imaging. Arthritis Rheum 56, 2986–2992. 10.1002/art.22851.

30. Li, Y., Xiao, W., Luo, W., Zeng, C., Deng, Z., Ren, W., Wu, G., and Lei, G. (2016). Alterations of amino acid metabolism in osteoarthritis: its implications for nutrition and health. Amino Acids 48, 907–914. 10.1007/s00726-015-2168-x.

31. Rajasekaran, S., Soundararajan, D.C.R., Tangavel, C., Muthurajan, R., Sri Vijay Anand, K.S., Matchado, M.S., Nayagam, S.M., Shetty, A.P., Kanna, R.M., and Dharmalingam, K. (2020). Human intervertebral discs harbour a unique microbiome and dysbiosis determines health and disease. Eur Spine J 29, 1621–1640. 10.1007/s00586-020-06446-z.

32. Durazzi, F., Sala, C., Castellani, G., Manfreda, G., Remondini, D., and De Cesare, A. (2021). Comparison between 16S rRNA and shotgun sequencing data for the taxonomic characterization of the gut microbiota. Sci Rep 11, 3030. 10.1038/s41598-021-82726-y.

33. Vázquez, L., Flórez, A.B., Redruello, B., and Mayo, B. (2020). Metabolism of Soy Isoflavones by Intestinal Bacteria: Genome Analysis of an Adlercreutzia Equolifaciens Strain That Does Not Produce Equol. Biomolecules 10. 10.3390/biom10060950.

34. Jenks, B.H., Iwashita, S., Nakagawa, Y., Ragland, K., Lee, J., Carson, W.H., Ueno, T., and Uchiyama, S. (2012). A pilot study on the effects of S-equol compared to soy isoflavones on menopausal hot flash frequency. J Womens Health (Larchmt) 21, 674–682. 10.1089/jwh.2011.3153.

35. Scanlan, P.D., Shanahan, F., and Marchesi, J.R. (2009). Culture-independent analysis of desulfovibrios in the human distal colon of healthy, colorectal cancer and polypectomized individuals. FEMS Microbiol Ecol 69, 213–221. 10.1111/j.1574-6941.2009.00709.x.

36. Distrutti, E. (2011). Hydrogen sulphide and pain. Inflamm Allergy Drug Targets 10, 123–132. 10.2174/187152811794776240.

37. Yazici, C., Wolf, P.G., Kim, H., Cross, T.L., Vermillion, K., Carroll, T., Augustus, G.J., Mutlu, E., Tussing-Humphreys, L., Braunschweig, C., et al. (2017). Race-dependent association of sulfidogenic bacteria with colorectal cancer. Gut 66, 1983–1994. 10.1136/gutjnl-2016-313321.

38. Engels, C., Ruscheweyh, H.J., Beerenwinkel, N., Lacroix, C., and Schwab, C. (2016). The Common Gut Microbe Eubacterium hallii also Contributes to Intestinal Propionate Formation. Front Microbiol 7, 713. 10.3389/fmicb.2016.00713.

39. Roth-Schulze, A.J., Penno, M.A.S., Ngui, K.M., Oakey, H., Bandala-Sanchez, E., Smith, A.D., Allnutt, T.R., Thomson, R.L., Vuillermin, P.J., Craig, M.E., et al. (2021). Type 1 diabetes in pregnancy is associated with distinct changes in the composition and function of the gut microbiome. Microbiome 9, 167. 10.1186/s40168-021-01104-y.

40. Grant, M.P., Epure, L.M., Bokhari, R., Roughley, P., Antoniou, J., and Mwale, F. (2016). Human cartilaginous endplate degeneration is induced by calcium and the extracellular calcium-sensing receptor in the intervertebral disc. Eur Cell Mater 32, 137–151. 10.22203/ecm.v032a09.

41. Henke, M.T., Kenny, D.J., Cassilly, C.D., Vlamakis, H., Xavier, R.J., and Clardy, J. (2019). Ruminococcus gnavus, a member of the human gut microbiome associated with Crohn’s disease, produces an inflammatory polysaccharide. Proc Natl Acad Sci U S A 116, 12672–12677. 10.1073/pnas.1904099116.

42. Bernstein, C.N., Blanchard, J.F., Rawsthorne, P., and Yu, N. (2001). The prevalence of extraintestinal diseases in inflammatory bowel disease: a population-based study. Am J Gastroenterol 96, 1116–1122. 10.1111/j.1572-0241.2001.03756.x.

43. Breban, M., Tap, J., Leboime, A., Said-Nahal, R., Langella, P., Chiocchia, G., Furet, J.P., and Sokol, H. (2017). Faecal microbiota study reveals specific dysbiosis in spondyloarthritis. Ann Rheum Dis 76, 1614–1622. 10.1136/annrheumdis-2016-211064.

44. Machiels, K., Sabino, J., Vandermosten, L., Joossens, M., Arijs, I., de Bruyn, M., Eeckhaut, V., Van Assche, G., Ferrante, M., Verhaegen, J., et al. (2017). Specific members of the predominant gut microbiota predict pouchitis following colectomy and IPAA in UC. Gut 66, 79–88. 10.1136/gutjnl-2015-309398.

45. Zeng, S.L., Li, S.Z., Xiao, P.T., Cai, Y.Y., Chu, C., Chen, B.Z., Li, P., Li, J., and Liu, E.H. (2020). Citrus polymethoxyflavones attenuate metabolic syndrome by regulating gut microbiome and amino acid metabolism. Sci Adv 6, eaax6208. 10.1126/sciadv.aax6208.

46. Bloomgarden, Z. (2018). Diabetes and branched-chain amino acids: What is the link? J Diabetes 10, 350–352. 10.1111/1753-0407.12645.

47. Tamanai-Shacoori, Z., Smida, I., Bousarghin, L., Loreal, O., Meuric, V., Fong, S.B., Bonnaure-Mallet, M., and Jolivet- Gougeon, A. (2017). Roseburia spp.: a marker of health? Future Microbiol 12, 157–170. 10.2217/fmb-2016-0130.

48. Machiels, K., Joossens, M., Sabino, J., De Preter, V., Arijs, I., Eeckhaut, V., Ballet, V., Claes, K., Van Immerseel, F., Verbeke, K., et al. (2014). A decrease of the butyrate- producing species Roseburia hominis and Faecalibacterium prausnitzii defines dysbiosis in patients with ulcerative colitis. Gut 63, 1275–1283. 10.1136/gutjnl-2013-304833.

49. Lim, C.K., Rideout, J.M., and Wright, D.J. (1983). High- performance liquid chromatography of naturally occurring 8-, 7-, 6-, 5- and 4-carboxylic porphyrin isomers. J Chromatogr 282, 629–641. 10.1016/s0021-9673(00)91640-6.

50. Borelli, C., Merk, K., Schaller, M., Jacob, K., Vogeser, M., Weindl, G., Berger, U., and Plewig, G. (2006). In vivo porphyrin production by P. acnes in untreated acne patients and its modulation by acne treatment. Acta Derm Venereol 86, 316–319. 10.2340/00015555-0088.

51. Wollenberg, M.S., Claesen, J., Escapa, I.F., Aldridge, K.L., Fischbach, M.A., and Lemon, K.P. (2014). Propionibacterium- produced coproporphyrin III induces Staphylococcus aureus aggregation and biofilm formation. mBio 5, e01286–01214. 10.1128/mBio.01286-14.

52. Aghazadeh, J., Salehpour, F., Ziaeii, E., Javanshir, N., Samadi, A., Sadeghi, J., Mirzaei, F., and Naseri Alavi, S.A. (2017). Modic changes in the adjacent vertebrae due to disc material infection with Propionibacterium acnes in patients with lumbar disc herniation. Eur Spine J 26, 3129–3134. 10.1007/s00586-016-4887-4.

53. Carricajo, A., Nuti, C., Aubert, E., Hatem, O., Fonsale, N., Mallaval, F.O., Vautrin, A.C., Brunon, J., and Aubert, G. (2007). Propionibacterium acnes contamination in lumbar disc surgery. J Hosp Infect 66, 275–277. 10.1016/j.jhin.2007.04.007.

54. Pittenger, M.F., Mackay, A.M., Beck, S.C., Jaiswal, R.K., Douglas, R., Mosca, J.D., Moorman, M.A., Simonetti, D.W., Craig, S., and Marshak, D.R. (1999). Multilineage potential of adult human mesenchymal stem cells. Science 284, 143–147. 10.1126/science.284.5411.143.

55. Bartel, D.P. (2004). MicroRNAs: genomics, biogenesis, mechanism, and function. Cell 116, 281–297. 10.1016/s0092-8674(04)00045-5.

56. Louis, P., and Flint, H.J. (2017). Formation of propionate and butyrate by the human colonic microbiota. Environ Microbiol 19, 29–41. 10.1111/1462-2920.13589.

57. Laurent, G., German, N.J., Saha, A.K., de Boer, V.C., Davies, M., Koves, T.R., Dephoure, N., Fischer, F., Boanca, G., Vaitheesvaran, B., et al. (2013). SIRT4 coordinates the balance between lipid synthesis and catabolism by repressing malonyl CoA decarboxylase. Mol Cell 50, 686–698. 10.1016/j.molcel.2013.05.012.

58. Anderson, K.A., Huynh, F.K., Fisher-Wellman, K., Stuart, J.D., Peterson, B.S., Douros, J.D., Wagner, G.R., Thompson, J.W., Madsen, A.S., Green, M.F., et al. (2017). SIRT4 Is a Lysine Deacylase that Controls Leucine Metabolism and Insulin Secretion. Cell Metab 25, 838–855.e815. 10.1016/j.cmet.2017.03.003.

59. Feldman, J.L., Dittenhafer-Reed, K.E., and Denu, J.M. (2012). Sirtuin catalysis and regulation. J Biol Chem 287, 42419–42427. 10.1074/jbc.R112.378877.

60. Segata, N., Izard, J., Waldron, L., Gevers, D., Miropolsky, L., Garrett, W.S., and Huttenhower, C. (2011). Metagenomic biomarker discovery and explanation. Genome Biol 12, R60. 10.1186/gb-2011-12-6-r60.

61. McDonald, D., Price, M.N., Goodrich, J., Nawrocki, E.P., DeSantis, T.Z., Probst, A., Andersen, G.L., Knight, R., and Hugenholtz, P. (2012). An improved Greengenes taxonomy with explicit ranks for ecological and evolutionary analyses of bacteria and archaea. Isme j 6, 610–618. 10.1038/ismej.2011.139.

62. Bolger, A.M., Lohse, M., and Usadel, B. (2014). Trimmomatic: a flexible trimmer for Illumina sequence data. Bioinformatics 30, 2114–2120. 10.1093/bioinformatics/btu170.

63. Quirici, N., Soligo, D., Bossolasco, P., Servida, F., Lumini, C., and Deliliers, G.L. (2002). Isolation of bone marrow mesenchymal stem cells by anti-nerve growth factor receptor antibodies. Experimental hematology 30, 783–791. 10.1016/s0301-472x(02)00812-3.

