## Supplemental Materials for "Disease-associated gut microbiome and metabolome changes in chronic low back pain patients with bone marrow lesions"

### Supplementary figures

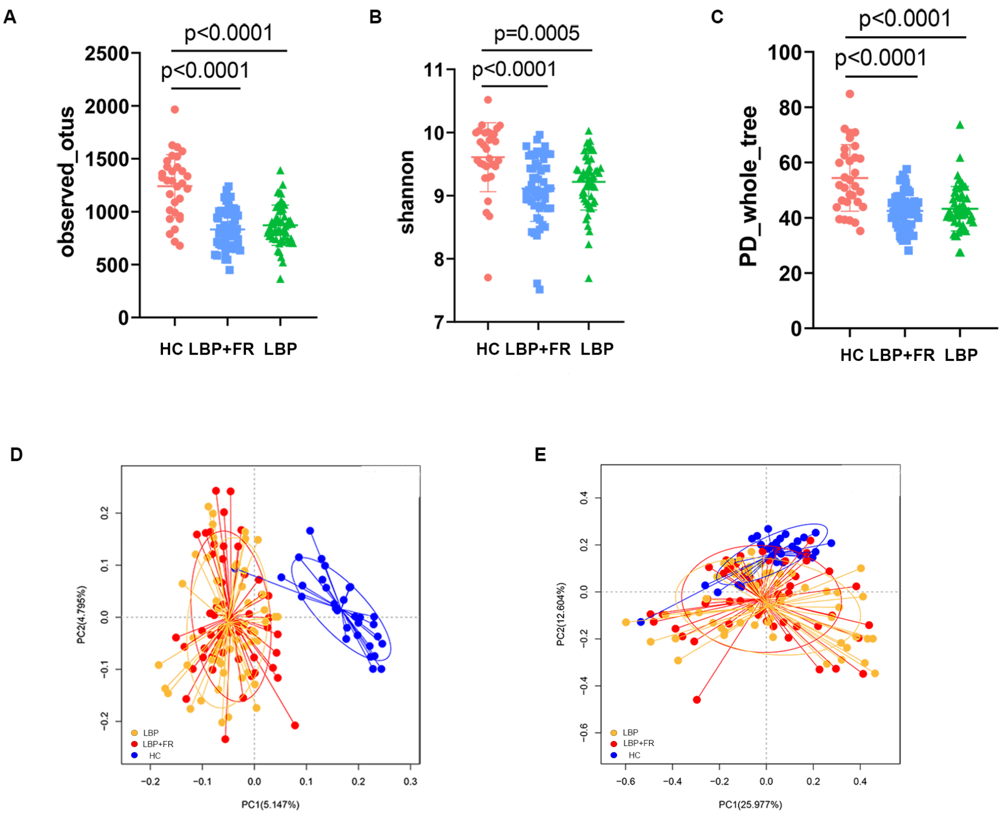


**Supplementary Figure 2. Alpha diversity indices and Beta diversity of LBP+FR, LBP and HC control stool samples**

(A-C) Alpha diversity based on Observed-otus, Shannon's diversity index and PD-whole-tree index of LBP+FR, LBP and HC faecal samples.

(D-E) Beta diversity based on Unweighted_UniFrac and Weighted_Unifrac distance of LBP+FR, LBP and HC faecal samples.

The sample size is n=31 HC, n=54 LBP+FR, n=53 LBP as biologically independent samples.

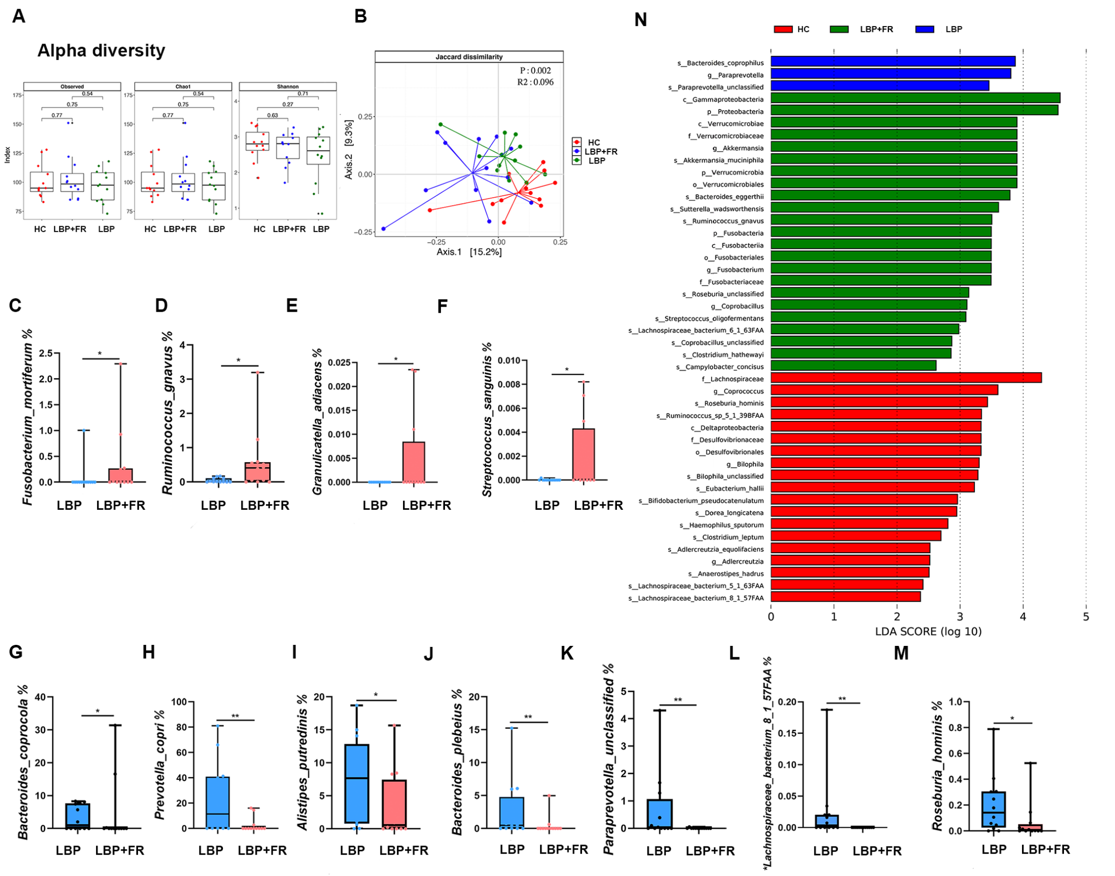


**Supplementary Figure 3. Differences in bacterial composition between HC, LBP+FR and LBP cohorts.**

(A) Comparison of alpha-diversity indices (Observed-otus, Chao-1-richness index and Shannon's diversity index) between LBP+FR, LBP and HC groups.

(B) Principal coordinate analysis (PCoA) based on Jaccard dissimilarity distances for bacterial sequences between LBP+FR, LBP and HC groups.

(C-M) The relative abundances of 11 genera showed significant differences at the species level between LBP+FR and LBP groups. * indicates $p$ <0.05; ** indicates $p$ <0.01; and *** indicates $p$ <0.001 by Wilcoxon rank-sum test.

(N) Linear discriminant analysis effect size (LEfSe) analysis identified different taxa between LBP+FR, LBP and HC groups. The LDA scores ($\log_{10}$) > 2 are listed.

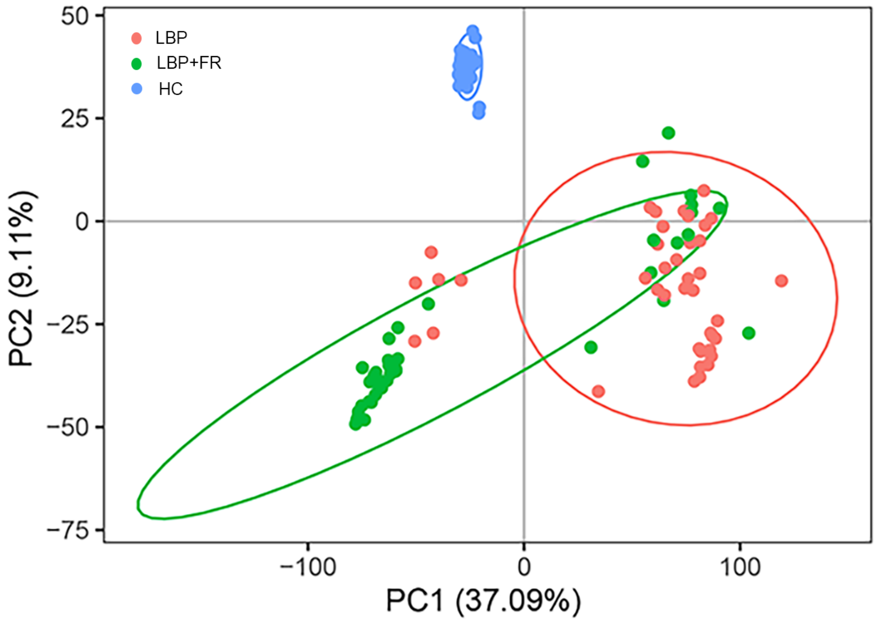


**Supplementary Figure 4. Differences in bacterial composition between HC, LBP+FR and LBP cohorts.**

The PCA score plot of serum metabolites in HC, LBP+FR and LBP groups.

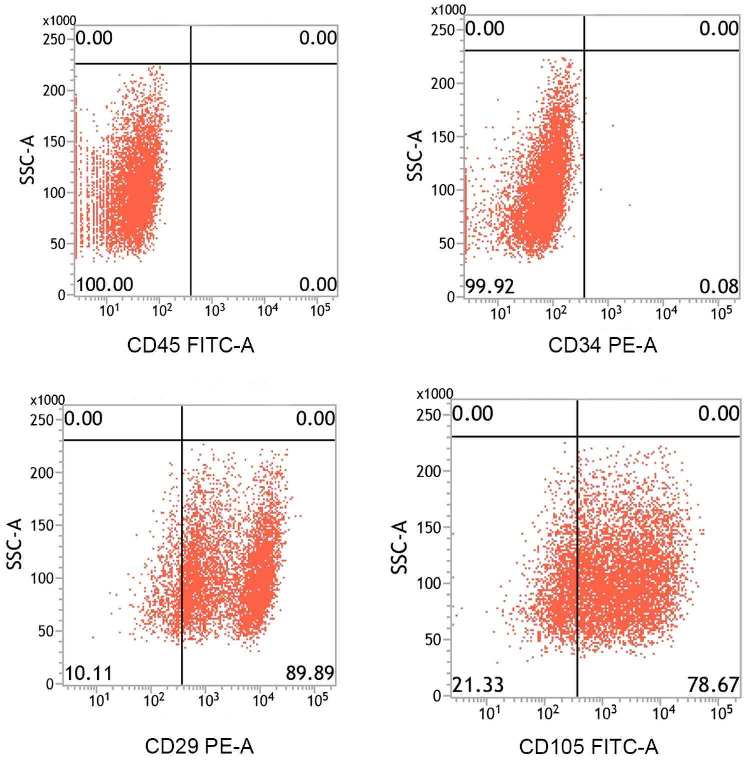


**Supplementary Figure 5. Flow identification of BM-MSCs.**

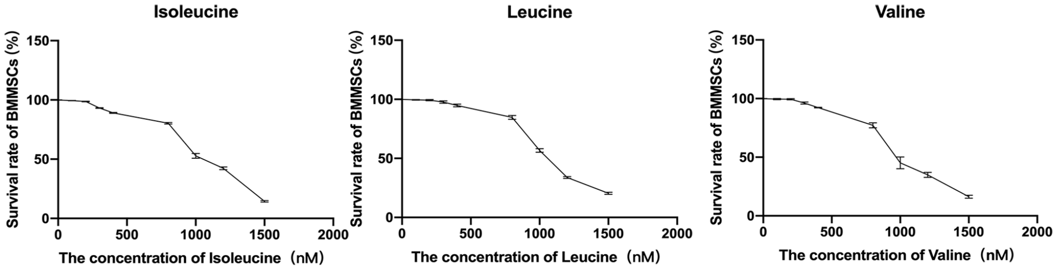


**Supplementary Figure 6. Increase in BCAA (valine, isoleucine, leucine) dose impacts on BM-MSCs viability in ex vivo model.**

Percentage of viable BM-MSCs, under the different stimulations with three kinds of BCAA demonstrating significant reduction in BM-MSCs viability (< 80%) at different doses. Valine, isoleucine and leucine were prepared to the concentration of 400 nM, 400 nM and 800 nM separately. Data are shown as mean (±SD). BM-MSCs viability was assessed by automated cell counters.

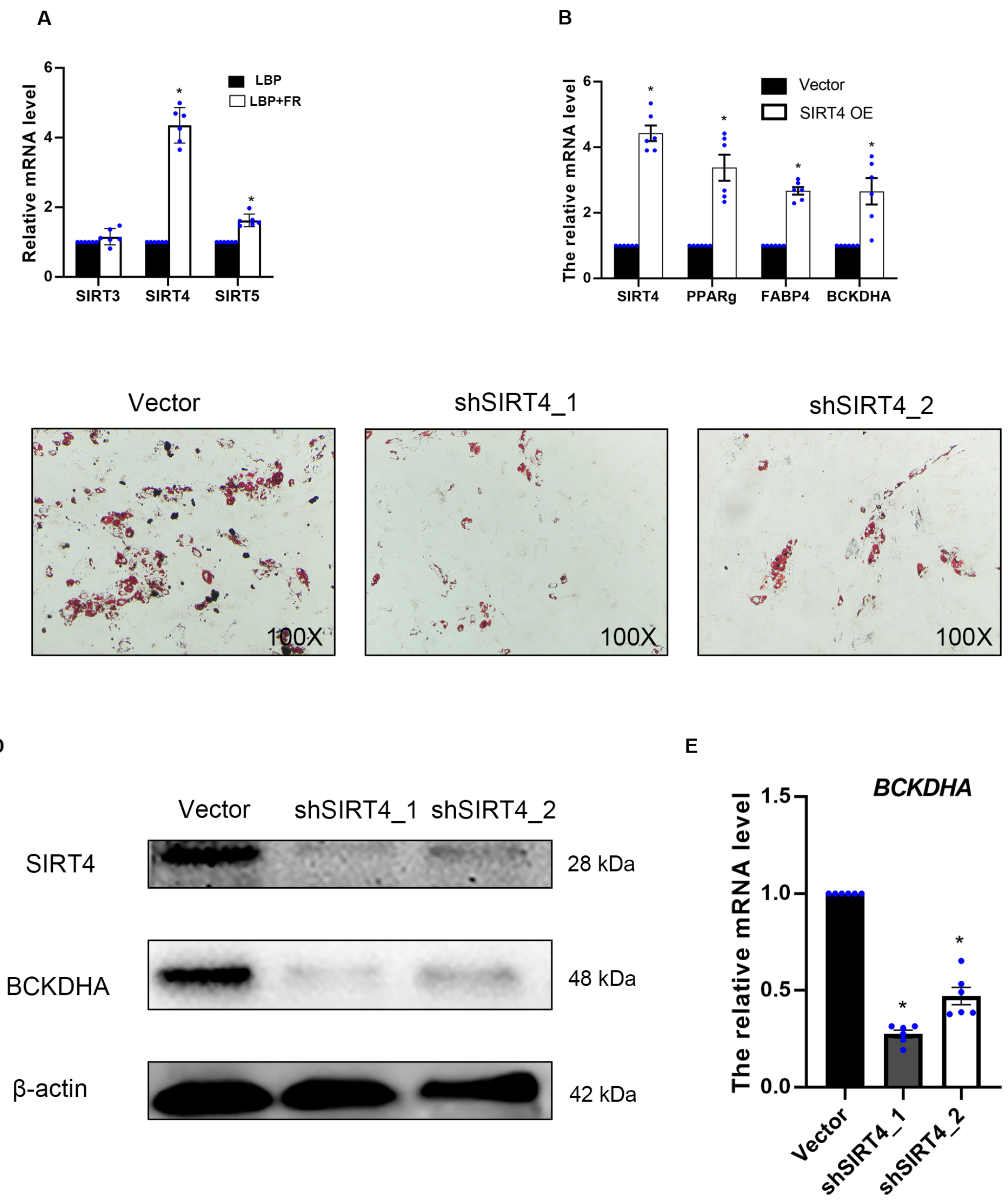


**Supplementary Figure 7. SIRT4 boosts adipogenesis on BM-MSCs.**

(A) RT-qPCR analysis results of SIRT3, SIRT4 and SIRT5 in BM-MSCs from LBP+FR and LBP group. (**P* < 0.05)

(B) qPCR gene expression analysis of SIRT4, PPARg, FABP4 and BCKDHA in control BM-MSCs or SIRT4-overexpressing BM-MSCs after 14 days of differentiation. (**P* < 0.05)

(C) Representative images of Oil Red O staining of lipids in shcontrol, shSIRT4_1 and shSIRT4_2 BM-MSCs differentiated for 14 days.100× magnification.

(D) Western blot analysis of SIRT4 and BCKDHA in shcontrol, shSIRT4_1 and shSIRT4_2 BM-MSCs differentiated for 14 days.

(E) RT-qPCR analysis results of BCKDHA in shcontrol, shSIRT4_1 and shSIRT4_2 BM-MSCs differentiated for 14 days. (**P* < 0.05)

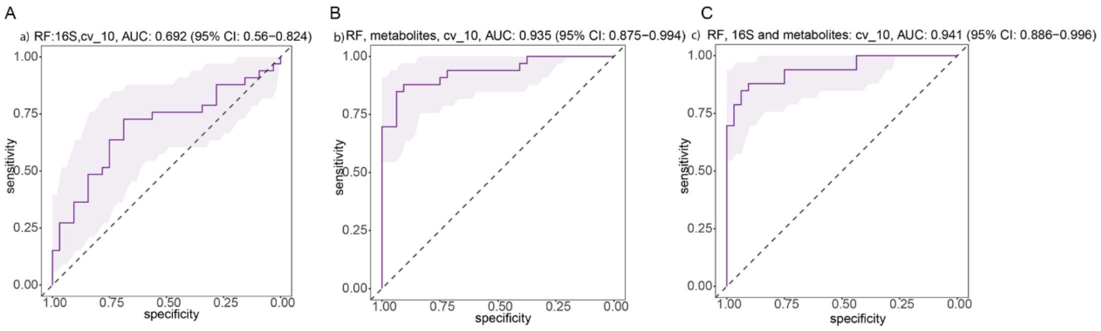


**Supplementary figure 8. Random Forest models to predict LBP different types.** Random Forest models discriminate the two groups with area under the curve (AUC) ranging from 0.69 to 0.94 (A) all 16S genus, AUC = 0.77; (B) all 10,000 metabolites, AUC = 0.935; (C) combination of all 16S genus and metabolites, AUC = 0.941.

**Supplementary Table 1.** A total of 343 discriminative bacterial species between HC, LBP+FR and LBP cohorts.

**Supplementary Table 2. Comparison of relative taxonomic abundance at family and genus level in HC, LBP+FR and LBP cohorts.**
Sample size is n=31 HC, n=54 LBP+FR, n=53 LBP as biologically independent samples. Data are shown as mean % ± standard error of the mean (SEM). Wilcoxon rank-sum Test calculated *P* values for 2 group comparisons. $P < 0.05$ considered statistically significant.

**Supplementary Table 3.** Comparison of underlying disease-correlated KEGG Orthologies (KOs) between LBP+FR, LBP and HC groups.

**Supplementary Table 4.** The top 50 differential fecal metabolites and enriched pathways in serum samples from the LBP+FR group.

**Supplementary Table 5.** RNA sequencing Detailed results of GO and KEGG enrichment analysis.

### References

1 Segata, N. *et al.* Metagenomic biomarker discovery and explanation. *Genome Biol* **12**, R60, doi:10.1186/gb-2011-12-6-r60 (2011).

2 McDonald, D. *et al.* An improved Greengenes taxonomy with explicit ranks for ecological and evolutionary analyses of bacteria and archaea. *Isme j* **6**, 610-618, doi:10.1038/ismej.2011.139 (2012).

3 Bolger, A. M., Lohse, M. & Usadel, B. Trimmomatic: a flexible trimmer for Illumina sequence data. *Bioinformatics* **30**, 2114-2120, doi:10.1093/bioinformatics/btu170 (2014).

4 Quirici, N. *et al.* Isolation of bone marrow mesenchymal stem cells by anti-nerve growth factor receptor antibodies. *Experimental hematology* **30**, 783-791, doi:10.1016/s0301-472x(02)00812-3 (2002).
